# Bcl-2 protein Noxa is required for metabolic reprogramming to glutamine dependence and for apoptosis in stimulated human CD8^+^ T cells

**DOI:** 10.64898/2025.12.08.692849

**Authors:** Tingyuan Yang, Himani Sharma, Jenna M. Benson, William J. Valente, Rebecca LaRue, Beau Webber, Branden Moriarity, Christopher A. Pennell, Stephen C. Jameson, Ameeta Kelekar

**Affiliations:** Department of Pharmacology, University of Minnesota, Minneapolis, MN 55455, USA; Department of Laboratory Medicine and Pathology, University of Minnesota, Minneapolis, MN 55455, USA; Center for Immunology, University of Minnesota, Minneapolis, MN 55455, USA; Minnesota Supercomputing Institute, University of Minnesota, Minneapolis, MN 55455, USA; Department of Pediatrics, University of Minnesota, Minneapolis, MN 55455, USA; Division of Pediatric Hematology and Oncology, University of Minnesota, Minneapolis, MN 55455, USA; Masonic Cancer Center, University of Minnesota, Minneapolis, MN 55455, USA

## Abstract

Naïve or memory T cells reprogram their metabolism upon antigenic stimulation. They increase their glucose uptake, relying on aerobic glycolysis for generating biomass while switching to glutamine to fuel energy production. Here we have identified a requirement for human Bcl-2 family, Noxa, in the metabolic switch to glutamine dependence in activated CD8^+^ T cells, that is independent of its canonical role in apoptosis at the end of the immune response. Using an in vitro co-stimulation model, we demonstrate that Noxa is induced in CD8^+^ T cells and remains elevated during the proliferative and differentiation phases of the response and through the onset of apoptosis. Noxa protein induction requires glutamine, is mediated via mTOR, and is independent of glutaminolysis. Glutamine, in turn, requires Noxa to facilitate its conversion to glutamate. CD8^+^ T cells lacking Noxa showed reduced levels of intracellular glutamate but no impairment of mitochondrial or effector function, and decreased dependence on glutamine for both respiration and growth during the proliferative phase. *NOXA* knockout CD8⁺ T cells also displayed significantly higher viability in the apoptotic phase of the immune response. CD8^+^ T cells from a human *NOXA* gene-replacement mouse responded normally to in vitro stimulation and in vivo acute infection. However, human Noxa-expressing murine CD8^+^ T cells displayed a distinctly proliferative gene signature in their transcriptome following activation, supporting an early growth-promoting role for this BH3-only protein. Our studies suggest that knocking out *NOXA* in human CD8^+^ T cells to increase their lifespan as well as their ability to survive and function in glutamine-poor microenvironments could be a promising immunotherapeutic strategy.

## INTRODUCTION

CD8⁺ cytotoxic T cells are central mediators of anti-tumor immunity and primary targets of multiple immunotherapeutic strategies. Naïve CD8^+^ T cells launch antigen-specific immune responses through a tightly regulated sequence of events—activation, clonal expansion, acquisition of cytotoxic effector functions, apoptosis, ending with memory formation—with each event coupled to distinct and dynamic metabolic states^1^. Upon antigenic stimulation, CD8⁺ T cells undergo rapid metabolic reprogramming, shifting from a quiescent state dependent on oxidative phosphorylation (OxPhos) to an anabolic state dominated by aerobic glycolysis^2^. This metabolic switch supports rapid proliferation and differentiation to effector cytotoxic T lymphocytes (CTLs) by supplying ATP, biosynthetic precursors, and signaling intermediates. The reprogramming diverts glucose for biosynthesis, while simultaneously increasing reliance on amino acids such as glutamine to sustain mitochondrial activity for homeostasis and energy production. In contrast, long-lived memory T cells adopt a metabolism that relies more on fatty acid oxidation (FAO), reflecting their reduced energetic demands and need for longevity^3, 4^.

Current immunotherapeutic strategies aimed at enhancing CD8⁺ T cell activation, cytotoxicity, and persistence, such as immune checkpoint blockade and adoptive T cell transfer, target both functional and metabolic programs. Their success is, however, often constrained by the hostile tumor microenvironment (TME), where nutrient deprivation and immunosuppressive signals can limit T cell expansion, survival, effector function and hinder memory cell formation. Thus, a better understanding of the metabolic regulation of T cell growth and expansion and the mechanisms underlying T cell survival and death is critical for optimizing these therapies and achieving durable tumor control.

Bcl-2 proteins are canonical apoptosis regulators, grouped into subfamilies based on shared Bcl-2 Homology (BH) domains^5, 6^. Enhancing the expression of pro-survival members, such as Bcl-2 and Bcl-x_L_, in chimeric antigen receptor (CAR) T cells has been shown to improve their expansion, persistence, and anti-tumor activity while reducing apoptotic priming and functional exhaustion^7^. In contrast, pro-apoptotic Bcl-2 proteins, particularly the BH3-only members, serve as critical checkpoints that initiate cell death under stress conditions^5^. Noxa, a phorbol-12-Myristate-13-Acetate-Induced Protein (PMAIP)^8^, is a BH3-only protein which promotes apoptosis by engaging the pro-survival protein Mcl-1 through its BH3 domain^9, 10^. Although human (h) Noxa was initially identified as a pro-apoptotic protein in T leukemia (T-ALL) cells, it was later shown to function as a growth promoter when stably phosphorylated on S13 in its N-terminus^11^. This phosphorylation, which is glucose-dependent, reduces Noxa’s structural flexibility and masks its BH3 domain, effectively silencing its apoptotic function by preventing a direct interaction with Mcl-1^12^. Dephosphorylation of the protein occurs following kinase inhibition or glucose deprivation, exposing the BH3 domain and triggering apoptosis^11^. Unexpectedly, however, Noxa also regulates cellular metabolism in T leukemia cells: its overexpression imparts a glucose-dependent Warburg-like proliferative phenotype, concomitantly increasing glutamine uptake and utilization for mitochondrial OxPhos^11, 13^. Additionally, Noxa is induced in primary human T cells following antigenic stimulation, suggesting an early role in a normal T cell immune response independent of apoptosis^11^.

It should be noted that hNoxa is the smallest of the Bcl-2 proteins, with 54 residues and a single BH3 domain (S1a). It is also significantly different from its murine homologue, mNoxa, which is 103 residues long, has two BH3 domains and lacks the regulatory serine in its N-terminal region^11, 14^. Moreover, mNoxa, like all other BH3-only Bcl-2 proteins, has been classified as a canonical apoptosis promoter, based on in vivo studies in knockout mice^15–17^. hNoxa’s involvement in both metabolic and pro-apoptotic regulation in T-ALL cells, and the importance of metabolic reprogramming in a T cell’s response to antigenic stimulation, led us to propose that the human protein promotes both proliferative metabolism and apoptosis in T cells responding to antigen. For the study described here, we established a basic in vitro model of the human CD8⁺T cell immune response to observe Noxa expression and interrogate Noxa’s role during the response. Our findings reveal that hNoxa contributes to the metabolic switch to glutamine dependence during early activation while also regulating apoptosis during the contraction phase.

## RESULTS

### Noxa protein is induced in stimulated human CD8^+^ T cells and remains highly expressed through the expansion and contraction phases of the immune response

We had previously determined that Noxa, was expressed at high levels in human T leukemia cells and in primary human T cells stimulated in vitro with anti-CD3/CD28 antibodies ^11^. Here we show both Noxa mRNA and protein are rapidly induced in human T cells upon activation, and that sustained stimulation is required for *NOXA* gene transcription (Fig. 1a, b). To further investigate the role of Noxa in the T cell immune response, we focused on the CD8^+^ T cell subset given their high proliferative capacity in response to antigen in short-term experiments. Moreover, a CD8^+^ T model would allow us to follow Noxa expression through proliferation, differentiation, and apoptosis, to memory cell formation, in a largely single subset in contrast to a CD4^+^ or pan-T cell model. Significant induction of both Noxa mRNA and protein was observed in naïve CD8^+^ T cells within 40 hours of co-stimulation in vitro (Fig. 1c, d). In contrast, Bim, another BH3 only member of the Bcl-2 family associated with T cell activation^16, 18^, was detected in naïve, unstimulated as well as stimulated T cell populations (S1b, c). Early activation of CD8^+^ T cells was supported by increase in CD69 and decrease in CD62L levels 40 hours post-stimulation prior to detectable cell differentiation or proliferation (Fig. 1e, f, S1d, e). The increased oxygen consumption rate (OCR) and spare respiratory capacity (SRC) of the CD8^+^ T cells are indicative of the metabolic shift to increased mitochondrial respiratory activity upon activation^19^ (Fig. 1g, h). Noxa is also detected in response to alternative activating stimuli, independent of TCR engagement, such as PMA/ionomycin or concanavalin A (S1f, g), indicating that induction of the protein was related to T cell activation.

**Figure 1.**
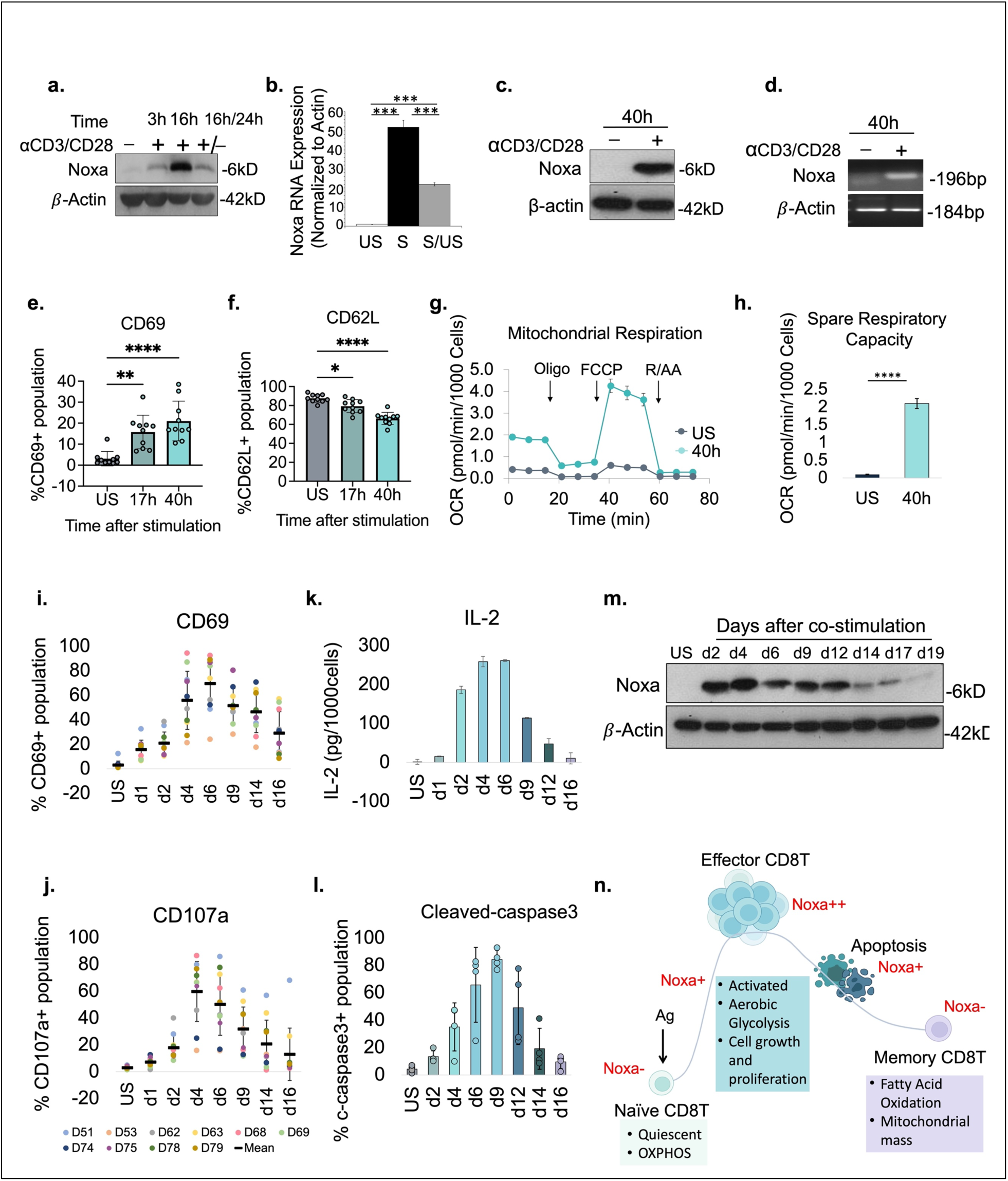
Bcl-2 protein Noxa is induced in human CD8^+^ T cells upon stimulation and remains detectable through expansion and contraction phases. **a.** Western blotting analysis of protein extracts from unstimulated, 3h and 16h stimulated pan T cells or cells stimulated for 16 hours followed by an additional 24-hour incubation period without activating antibodies. **b**. qPCR analysis of Noxa mRNA expression in the T cells from the unstimulated (US) and 24h stimulated (S) and 24+/24-(S/US) samples above. **c-d**. Western blotting analysis of protein extracts (**c**) and RT-PCR analysis (**d**) for the expression of Noxa mRNA from unstimulated and 40h stimulated CD8+ T cells (40mg protein loaded per lane in western blots unless otherwise stated). **e-h.** Levels of CD69 (**e**) and CD62L (**f**) on naïve unstimulated, and 17 and 40h co-stimulated CD8^+^ T cells (n=10). Oxygen consumption rate (OCR) (**g**) and spare respiratory capacity (SCR) (**h**) of resting and 40h co-stimulated naïve CD8^+^ T cells. Oligo., oligomycin; R/AA, rotenone/antimycin (n=5). **i-l.** Naïve CD8^+^ T cells were analyzed for CD69 (**i**), CD107a (**j**), IL-2 production (**k**), and cleaved caspase-3 levels (**l**) at indicated times after co-stimulation. Data represent 4-10 independent donors (D). **m.** Western blot analysis of Noxa protein expression at the indicated times in naïve CD8^+^ T cells following in vitro co-stimulation. **n**. Schematic representation of phases in a naïve CD8^+^ T cell immune response including the associated observed Noxa expression pattern.

We next tracked the CD8^+^ T response to co-stimulation in vitro for several days, assessing cells for activation, proliferation, differentiation and apoptosis at regular intervals after the initial exposure to activating antibodies. Typically, increase in CD69 and reduction in CD62L levels were detectable in naïve human CD8^+^ T within 24 hours of activation (Fig. 1i, S1h), while differentiation to effectors, indicated by increased degranulation marker CD107a, Granzyme B levels and IL-2 production, peaked at day 4-5 (Fig. 1j, k, S1i). Cells shifted to the contraction or apoptotic phase after 7 days following the initial stimulation, (Fig. 1l), with viable cells that persisted beyond this phase transitioning to a quiescent state around day 15-17 of stimulation (data not shown). Similar patterns were observed in total CD8^+^ T cell populations with smaller naïve cell compartments, upon co-stimulation in vitro (S1j-l). Noxa was consistently induced in human CD8^+^ T cells upon co-stimulation, and remained expressed at high levels through their growth, differentiation, and apoptotic phases. However, the protein was undetectable in the viable population that remained after the contraction phase (Fig. 1m, S1m). The model shown in Figure 1n broadly correlates observed Noxa expression levels with the phases in a CD8^+^ T cell immune response. The pattern of expression suggests an additional early role for human Noxa in stimulated CD8^+^ T cells beyond its canonical role in apoptosis.

### Memory-like CD8^+^ T cells lack Noxa expression and display altered fuel dependence

To determine whether the small population of viable cells remaining after the contraction phase was a memory-like subset, we tracked the stimulated naïve CD8^+^ T cells with additional markers of differentiation. Loss of naïve cell marker, CD45RA, coincided with a concomitant increase in CD45RO levels (Fig. 2a, S2a) and emergence of effector populations. Markers CD69, CD107α, IL-2 and cleaved caspase3 (Fig. 1i-l) peaked between d5 and d9 following stimulation, subsequently decreasing, by d17, to levels comparable to those in unstimulated CD8^+^ T cells, suggesting the T cells had transitioned to a quiescent, memory-like state. Although CD45RA and CD45RO have historically served to distinguish between naïve and memory T cell population, not all CD45RO^+^ cells are memory T cells^20–22^. We observed a consistent, significant drop in the late differentiation marker HLA-DR^23, 24^ in the post-apoptotic phase of the immune response, that marked the onset of the memory phase (Fig. 2b, S2b). Restimulation of the memory-like population triggered a rapid increase in CD69 and CD107a surface expression, concomitant decrease in CD62L (Fig. 2c, d, S2c), and early and significant induction of Noxa (Fig. 2e) compared to naïve populations, indicating the cells were primed for a rapid recall response in keeping with the memory cell phenotype. Additionally, CD45RO levels remained high and no CD45RA^+^ terminally differentiated effector memory cells (TEMRA) were detected (Fig. S2d), in the unstimulated or re-stimulated population.

**Figure 2.**
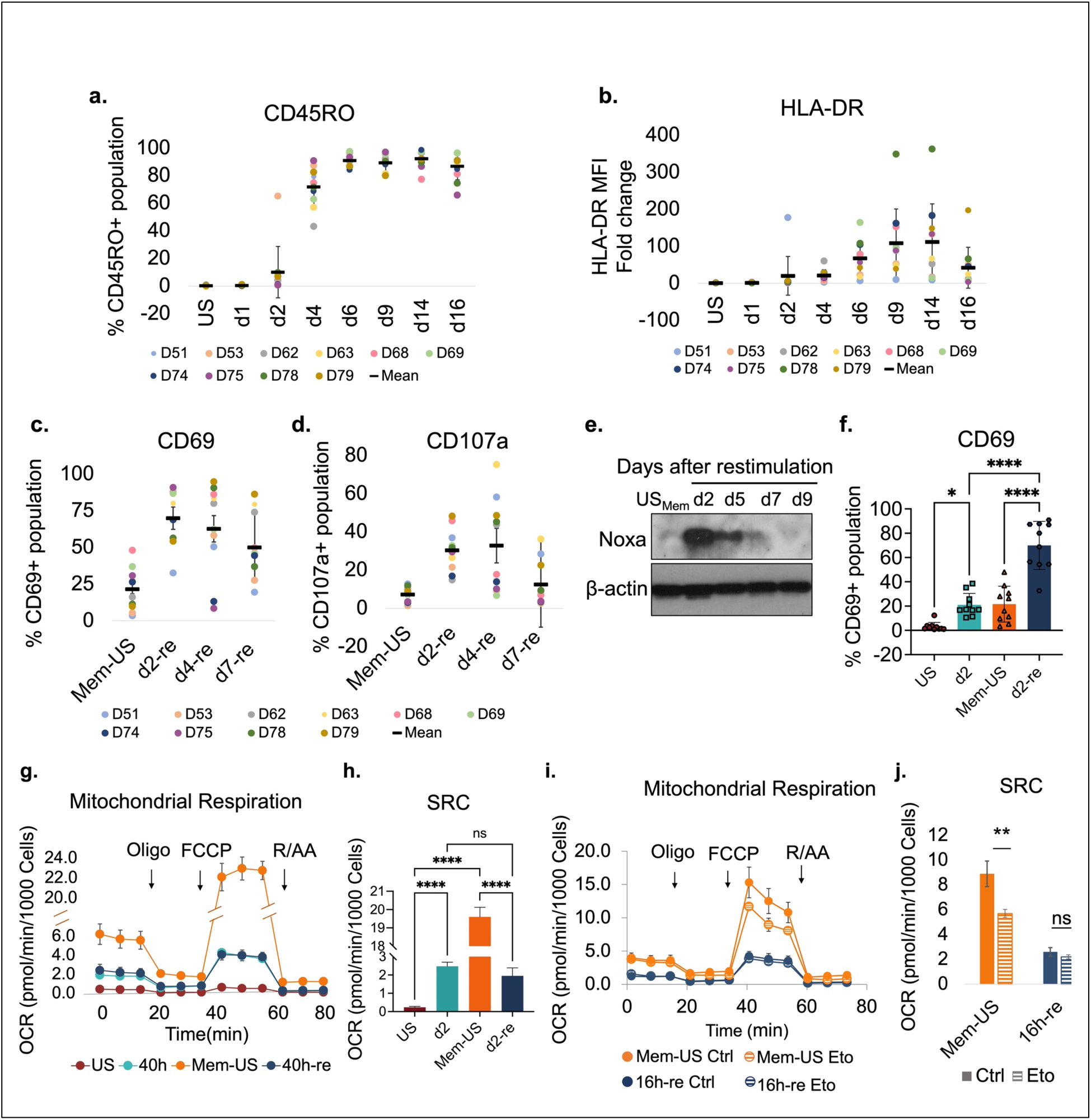
Memory CD8^+^ T cells at the end of the contraction phase lack Noxa expression and rely on fatty acid oxidation for mitochondrial respiration. **a-b.** Detection of percent CD45RO^+^ cells (**a**) and HLA-DR mean fluorescence intensity fold change relative to unstimulated (US) cells (**b)** at the indicated time points after stimulation of naïve CD8^+^ T cells (data are representative of 10 independent experiments) (D=Donor). **c-d.** Memory populations were re-stimulated and analyzed for CD69 (**c**) and CD107a (**d**) positivity. **e**. Representative western blots of the memory-like population showing Noxa expression at the indicated time points before and after re-stimulation. β-Actin was used as a loading control. **f.** Induction of CD69 in naïve and memory cells from matched donors 48h following co-stimulation (n=10). Data in **c**, **d** and **f** represent 4-10 independent experiments. **g-h.** Representative OCR (**g**) and spare respiratory capacity (SRC) (**h**) of naïve human CD8^+^ T cells in resting state, 40h after primary stimulation, unstimulated memory and memory cells 40h after re-stimulation. Oligo., oligomycin; R/AA, rotenone/antimycin. n=6 technical replicates. **i, j.** Representative OCR (**i**) and SRC (j) of memory CD8⁺ T cells 40 hours post-recall stimulation with or without etomoxir (Eto, 5 µM). Oligo., oligomycin; R/AA, rotenone/antimycin A. Data represent n = 6 technical replicates (n=5).

The memory-like unstimulated (Mem-US) cells also exhibited a metabolic profile consistent with the memory phenotype ^4, 25, 26^. In a Seahorse Mito Stress Test the Mem-US population displayed a significantly higher spare respiratory capacity than the re-stimulated memory-like as well as naïve co-stimulated populations from the same donor (Fig. 2 g, h). Memory CD8^+^ T cells preferentially utilize fatty acids as their primary fuel source, an adaptation that supports their long-term survival and rapid response upon re-encountering antigen ^25–27^. The reduced OCR and SRC on the Mem-US population in the presence of Etomoxir, inhibitor of carnitine palmitoyltransferase-1 (CPT1), a key enzyme in fatty acid oxidation (FAO), indicated a dependence on FAO typical of CD8^+^ T memory subsets. The FAO dependence was lost when cells were re-stimulated (Fig. 2i, j). Again, Noxa induction correlated with reactivation of cells and with the proliferative state.

### Noxa protein induction requires glutamine and is mediated via the mTOR pathway

Noxa has non-canonical growth and survival roles in T-ALL cells, imparting a Warburg-like phenotype and enhancing uptake of both glucose and glutamine in proliferating populations ^11, 13^. Normal human T cells, likewise, switch to aerobic glycolysis and utilize glutamine as a source of fuel for the energy-producing mitochondrial TCA cycle as they transition to a proliferative phase in response to antigen exposure ^28, 29^. Mitochondrial oxygen consumption, spare respiratory capacity and proliferation are, correspondingly, impaired significantly in 24h stimulated CD8^+^ T cells in the absence of glutamine (Fig. 3a, b, S3a). Notably, activated CD8^+^ T cells failed to induce Noxa protein in glutamine-free (G-F) medium; this regulation is at the level of translation as Noxa mRNA induction is not suppressed and, often, even enhanced, in G-F medium (Fig. 3c, d).

**Figure 3.**
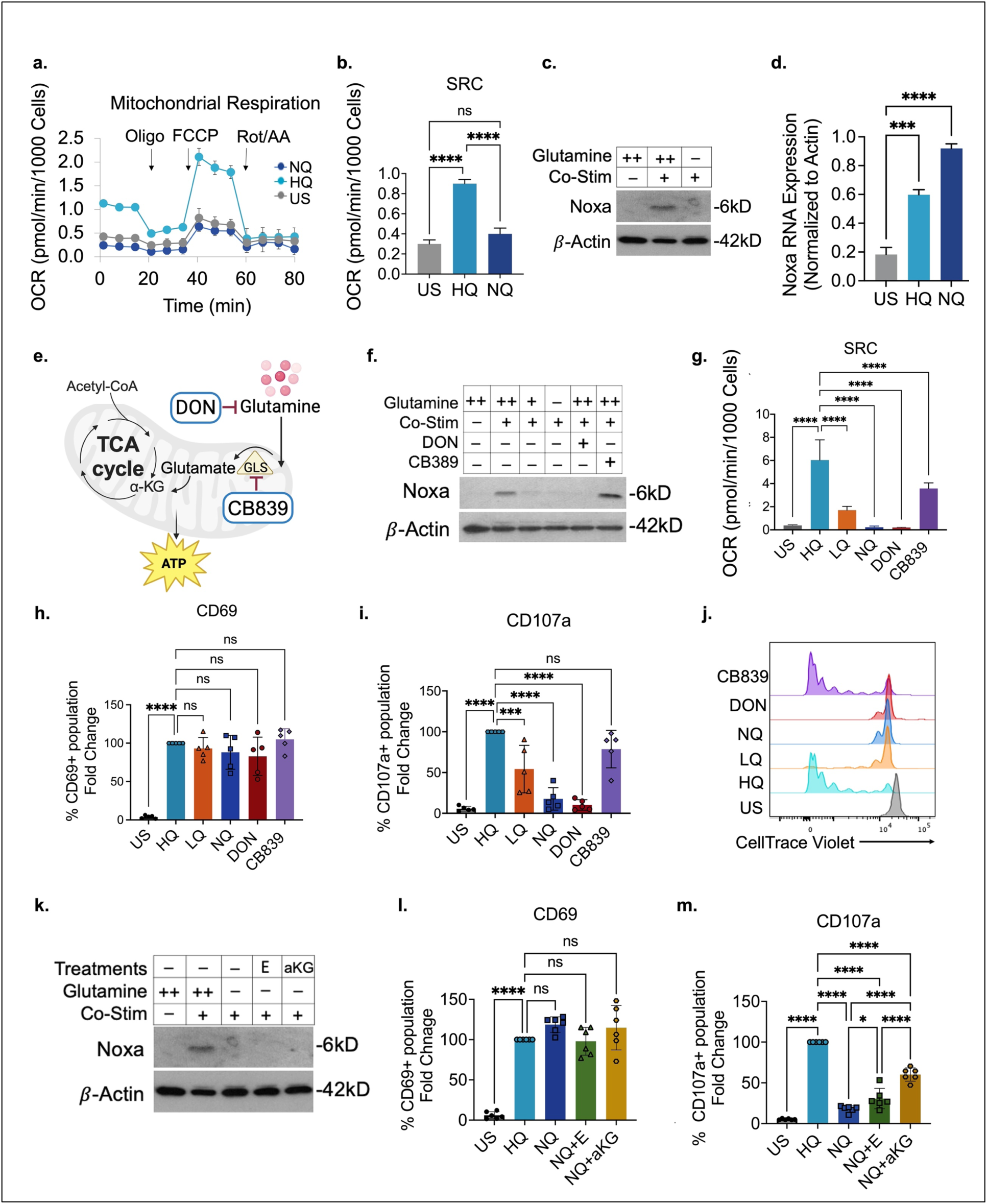
Noxa protein induction requires glutamine. **a-b.** Mito Stress Seahorse assay showing oxygen consumption rate (OCR) and spare respiratory capacity (SRC) of unstimulated (US) and 24-hour co-stimulated human naïve CD8⁺ T cells cultured in the presence (HQ) or absence (NQ) of glutamine (n = 6). **c-d.** Western blot and RT-qPCR analysis of Noxa protein and mRNA levels in CD8⁺ T cells 40 h after stimulation with anti-CD3/CD28 antibodies in high glutamine or glutamine-free medium. **e.** Cartoon depicting entry of glutamine derived metabolites into the mitochondrial TCA cycle and points of action of glutaminase (GLS) inhibitor, CB839, and glutamine antagonist, 6-Diazo-5-oxo-L-norleucine, DON. **f.** Noxa expression in CD8⁺ T cells cultured in HQ (high glutamine, 2mM), LQ (low glutamine, 0.2mM), NQ (No glutamine, 0mM), or in the presence of inhibitors CB839 (2 µM) or DON (200 µM) for 40 h. **g-j.** Evaluation of the same cell populations under these conditions for spare respiratory capacity, SRC (**g**) and percent CD69⁺ cells (**h**) on day 2, percent CD107a⁺ cells (**i**) and proliferation profiles (**j**) by day 4, following TCR engagement (n = 8). **k-m.** CD8^+^ T cells were co-stimulated in HQ, NQ medium, or (NQ) medium supplemented with cell permeable glutamine-derived metabolites dimethyl (DM)-glutamate (E) or DM-α-ketoglutarate (αKG) and evaluated for Noxa expression 40h later (**k**), for CD69^+^ activation on day 2 (**l**), and for CD107a⁺ degranulation on day 4 (**m**) (n = 6).

To determine whether glutaminolysis was essential for the observed induction of Noxa we turned to two inhibitors - glutaminase (GLS) inhibitor CB-839 and glutamine antagonist, 6-Diazo-5-oxo-L-norleucine or DON ^30^ (Fig. 3e). CB-839 had no effect on Noxa protein induction but DON, a broad-based inactivator of glutamine-utilizing reactions, abolished it. Blocking glutamine function or reducing its availability during receptor engagement had no impact on early activation but DON exerted effects similar those of glutamine deprivation on mitochondrial fitness, proliferation, differentiation and late activation of the CD8^+^ T cells (Fig. 3f-j, S3b-e). Additionally, replacing glutamine with cell-permeable versions of glutamate (E) or α-ketoglutarate (α-KG), did not restore Noxa expression but partially rescued CD8^+^ T cell cytotoxic potential (Fig. 3e, k-m). Together the data show that Noxa protein translation in activated CD8^+^ T cells is tied to the availability of glutamine rather than to its utilization via glutaminolysis.

Given the pivotal role of glutamine in the activation of mTORC1^31, 32^, we hypothesized that glutamine was modulating Noxa translation indirectly, via the mTORC1 pathway. mTORC1 promotes protein translation through effector molecules, eukaryotic initiation factor binding protein 4EBP-1 and ribosomal protein S6K^33, 34^. Inhibition of mTORC1 activity with rapamycin and everolimus^35^ in the presence of glutamine, recapitulated the phenotype observed under glutamine deprivation, suppressing Noxa induction in stimulated CD8^+^ T cells without affecting early activation (Fig. 4a, S4a). Similarly, both glutamine deprivation and pharmacological mTORC1 inhibition reduced phosphorylation of 4E-BP1 and S6, target of S6K, and impaired differentiation and proliferation of the T cells (Fig. 4b-d, S4b).

**Figure 4.**
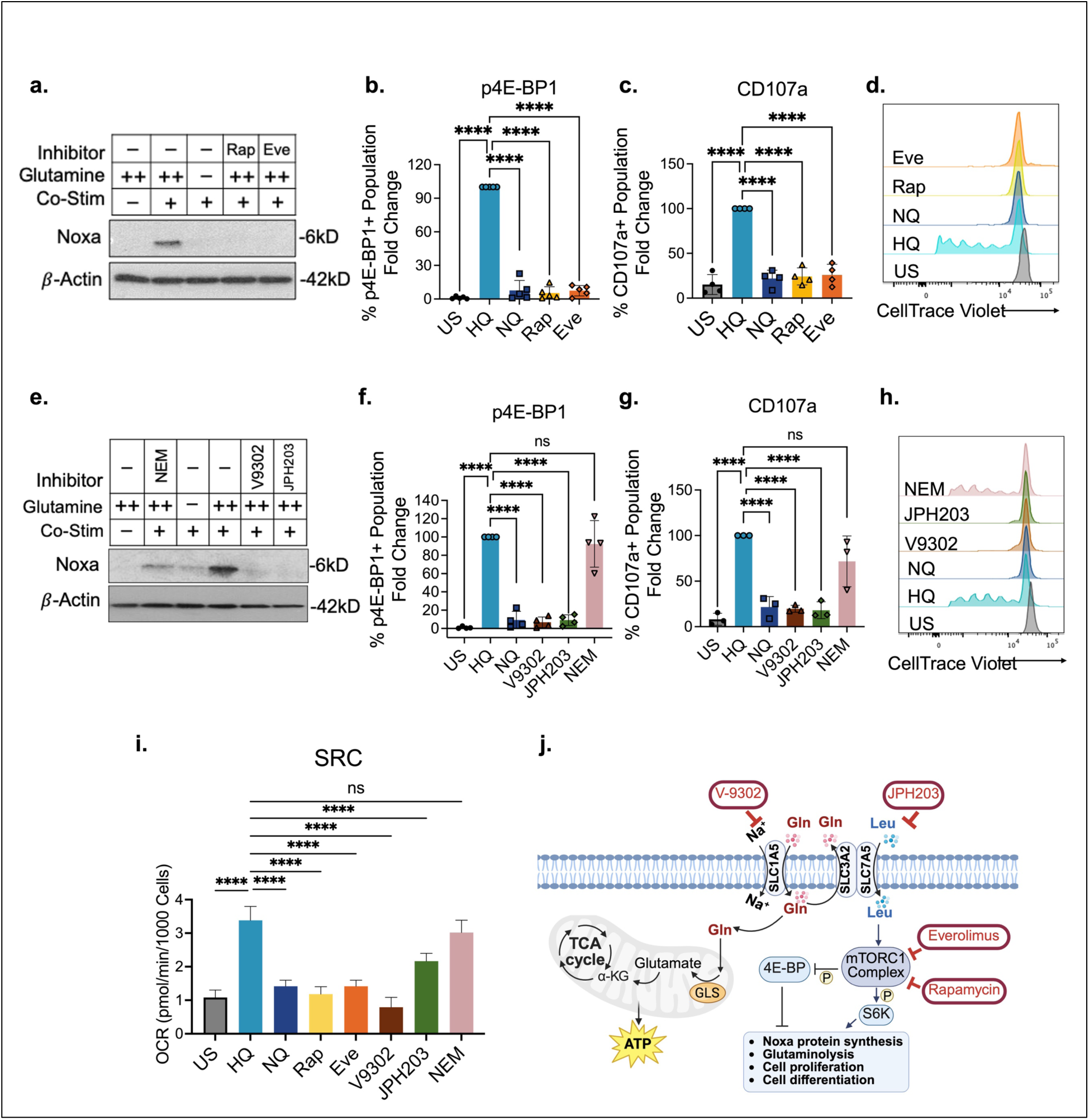
Noxa protein induction requires glutamine and is mediated via the mTORC1 pathway. **a.** Noxa protein expression in human CD8⁺ T cells 48h after TCR co-stimulation with or without glutamine (2uM), or in the presence of mTORC1 inhibitors 100nM rapamycin (Rap) or 100nM everolimus (Eve). **b-d.** Evaluation of phospho-4E-BP1 (**b**) and CD107a (**c**) expression (n=4-5, data shown as fold change relative to HQ group) and proliferation of CD8^+^ T cells (**d**) in the presence of the inhibitors. CTV profiles are from a representative donor. **e.** Noxa levels in CD8^+^ T co-stimulated for 48h with or without glutamine (2uM), or in the presence of inhibitors V9302 (2uM), JPH203 (1uM) or NEM (100 µM)**. f-h**. Levels of p4E-BP1 (**f**) and CD107a (**g**) in treated CD8^+^ T cells compared to HQ controls on day 4 of stimulation (n=4) and CTV profiles (**h**) from a representative donor**. i**. Mitochondrial SRC of unstimulated (US) and 40-hour post-stimulated CD8⁺ T cell cultured with or without glutamine, or treated with mTORC1 inhibitors, rapamycin and everolimus, or amino acid transporter inhibitors, V-9302, JPH203, and NEM. Abbreviations: Rap (rapamycin), Eve (everolimus), V-9302 (glutamine transporter SLC1A5 inhibitor), JPH203 (glutamine-leucine antiporter SLC3A2/SCL7A5 inhibitor) and NEM (CAT-1 inhibitor, N-ethylmaleimide). (n = 5). **j.** Cartoon illustrating showing how mTORC1 inhibitors affect major targets of mTORC1 activity and inhibit translation, and how glutamine controls Noxa protein levels via the mTORC1 pathway in activated CD8^+^ T cells.

Glutamine transporters and antiporters, such as ASCT2/SLC1A5, and SLC3A2/SLC7A5 and others, facilitate its uptake and/or rapid efflux in exchange for essential amino acids, such as leucine or cysteine, that are important for mTOR activation^32, 36^. Glutamine-leucine exchange is a rate-limiting step in mTORC1 activation. To determine the contribution of the mTORC1 activation to Noxa translation, we inhibited glutamine uptake with V-9302, blocked glutamine-leucine exchange with JPH203, and included N-ethylmaleimide (NEM) to inhibit other cysteine-dependent cationic (CAT) amino acid transporters^37–39^. Interfering with glutamine influx or glutamine-leucine exchange abrogated Noxa induction in stimulated CD8^+^ T cells while inhibition of CAT-1 by NEM was only weakly effective (Fig. 4e). Inactivation of mTORC1 signaling via glutamine perturbation was confirmed by the reduction in p4E-BP1 levels and led to impaired differentiation and proliferation, although T cell early activation remained unaffected (Fig. 4f-h and S4c). Mitochondrial respiratory capacity was significantly reduced in the presence of mTORC1 inhibition or glutamine transport blockade (Fig. 4i and S4d). Previous studies from other groups have shown that rapamycin treatment or genetic ablation of mTORC1 activator proteins lead to a reduction in the oxygen consumption rate of 24h stimulated murine CD4⁺ and CD8⁺ T cells, supporting our observations, and underscoring the role of mTORC1 in sustaining mitochondrial function during early T cell activation ^40, 41^.

Overall, these observations suggest that, while the activation of CD8^+^ T cells is independent of both glutamine and the mTOR pathway, glutamine controls Noxa protein synthesis in the activated cells through regulation of mTOR function. The model in Figure 4j summarizes the data from Figures 3. S3, 4 and S4.

### Noxa regulates both the metabolic reprogramming to glutamine dependence of CD8^+^ T cells and their entry into the apoptotic phase

We next asked if Noxa knockdown affected glutamine metabolism and mitochondrial health in activated CD8^+^ T cells. This was based on our observations that glutamine was essential both for Noxa induction and mitochondrial respiratory function in activated CD8^+^ T cells (Fig. 3c, f, Fig. 4j), and on previous studies in T leukemia cells showing that Noxa over-expression increased glutamine uptake and utilization in the TCA cycle^13^. Glutaminase inhibition by CB-839, used as a positive control, decreased glutamate production decisively in T cells within 48h of stimulation, without affecting Noxa expression. Suppression of Noxa with siRNA also led to significant reduction in intracellular glutamate levels suggesting that Noxa was, directly or indirectly, regulating the conversion of glutamine to glutamate (Fig. 5a-c). Noxa knockdown had no impact on CD8^+^ T cell activation or differentiation (Fig. 5d, S5a) while mitochondrial respiratory capacity and IL-2 production were modestly enhanced (Fig. 5e-g). This could indicate that partial blocking of glutaminolysis causes activated cells to rely on alternative ‘anaplerotic’ substrates to fuel the TCA cycle^42^.

**Figure 5.**
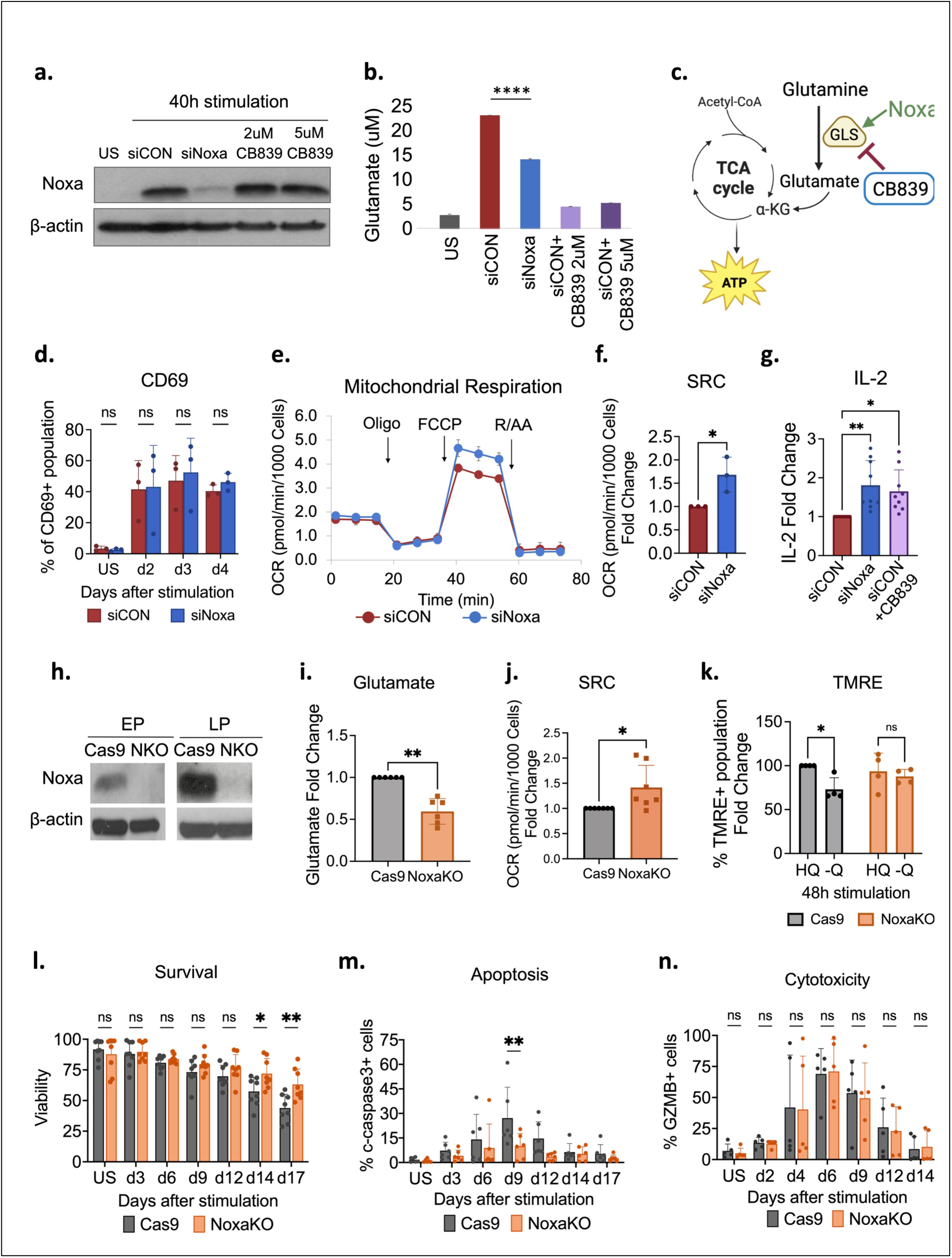
Noxa facilitates the switch to glutamine dependence and the entry into apoptosis of stimulated CD8^+^ T cells. **a, b.** Noxa protein level **(a)** and intracellular glutamate concentration **(b)** in unstimulated CD8^+^ T cells, 48h stimulated control (siCON) and Noxa-silenced (siNoxa) CD8^+^ T cells and siCON cells treated with 2uM and 5uM CB-839 (n=3). **c.** Schematic showing Noxa how regulates the entry of glutamine into the TCA cycle in activated T cells. **d.** Percent CD69⁺ cells in siCON- and siNoxa-treated CD8⁺ T cells at indicated time points following stimulation (n = 3). **e-g.** Metabolic analysis of siCON and siNoxa CD8^+^ T cells 48 hours post-stimulation, showing (**e**) oxygen consumption rate (OCR) and (**f**) spare respiratory capacity (SRC) (n=3). **g.** Fold change in IL-2 production in 48h stimulated siNoxa and 2uM CB-839-treated siCON CD8^+^ T cells relative to siCON controls (n=9). **h-j.** Noxa protein detection in CRISPR/Cas9 Noxa KO and Cas9-control CD8^+^ T cells, early passage (EP) and late passage (LP) CD8^+^ T cells, 48 hours after stimulation (**h**), and glutamate concentration (**i**), and spare respiratory capacity (**j**) in 48-hour stimulated Cas9-control and Noxa KO CD8^+^ T cells, expressed as fold change relative to Cas9-control group (n=6). **k.** TMRE fluorescence in Noxa KO cells activated for 48 hours in glutamine-rich (HQ) or glutamine-free (-Q) medium relative to Cas9-controls (n=4). **l-n**. Cell viability (**l**), percent cleaved-caspase-3 (**m**) and GZMB (**n**) positivity in Cas9-control and Noxa KO CD8^+^ T populations assessed by flow cytometry at indicated times post-stimulation (n=5-8).

Thus, while our data showed that induction of Noxa in CD8^+^ T cells is glutamine dependent (Fig. 3c, f, Fig. 4a), siRNA knockdown studies, which offer a window into the immediate effects of interfering with its induction, indicate Noxa may itself be required for optimal glutaminolysis (Fig. 5b). To assess the effects of Noxa loss at later times in our in vitro stimulation model, we generated *NOXA* knockout (NoxaKO) populations using CRISPR/Cas9 (Fig. 5h). Knockout cells showed enhanced mitochondrial fitness and reduced dependence on glutamine for respiration compared to Cas9 controls in the early activation phase. NoxaKO CD8⁺ T cells expanded in glutamine-replete media showed markedly reduced glutamate levels and increased mitochondrial spare respiratory capacity (Fig. 5i, j, S5b). NoxaKO cells were resistant to glutamine deprivation as indicated by TMRE labeling of cells stimulated for 24 hours in the absence of glutamine (Fig. 5k). Furthermore, Noxa deletion conferred a distinct survival advantage to CD8⁺ T cells overall, imparting increased cell viability and persistence and decreasing apoptotic cell death (Fig. 5l, m) without affecting activation, degranulation and, importantly, their cytotoxic potential, as indicated by Granzyme B production (S5c, d, Fig. 5n).

### CD8^+^ T lymphocytes from gene replacement mouse expressing hNoxa and wildtype mice respond similarly to antigenic stimulation

Our prior studies of human T leukemia cells had suggested that the survival function of human Noxa was largely attributable to the amino terminal modifiable serine^11, 12^ (S1a). The serine, notably absent in the murine protein (S6a), may also contribute to the extended growth phase observed in primary human T cells, at least in vitro, although this remains to be determined. We created, in collaboration with Biocytogen, a gene replacement (GR) h*NOXA* mouse, replacing the entire mouse *Pmaip 1* gene/regulatory region with the corresponding *Pmaip 1* sequence containing the gene and promoter for hNoxa (Fig. 6a). Specific reverse primers Mut-R and WT-R combined with a wild-type forward primer (WT-F), that amplify 399 bp or a 629 bp products in the GR and wildtype mice, respectively, are routinely employed for genotyping during breeding and confirming homozygous expression of the h*NOXA* allele in our GR mouse line (Fig. 6b, c). This model will enable us to investigate expression patterns and multiple functions of the human Noxa in both immune and non-immune tissue in the long term. The immediate goal relevant to this study, however, was to determine whether CD8^+^ T cells expressing the human protein were altered in any way in their response to antigenic stimulation in vitro or in vivo.

**Figure 6.**
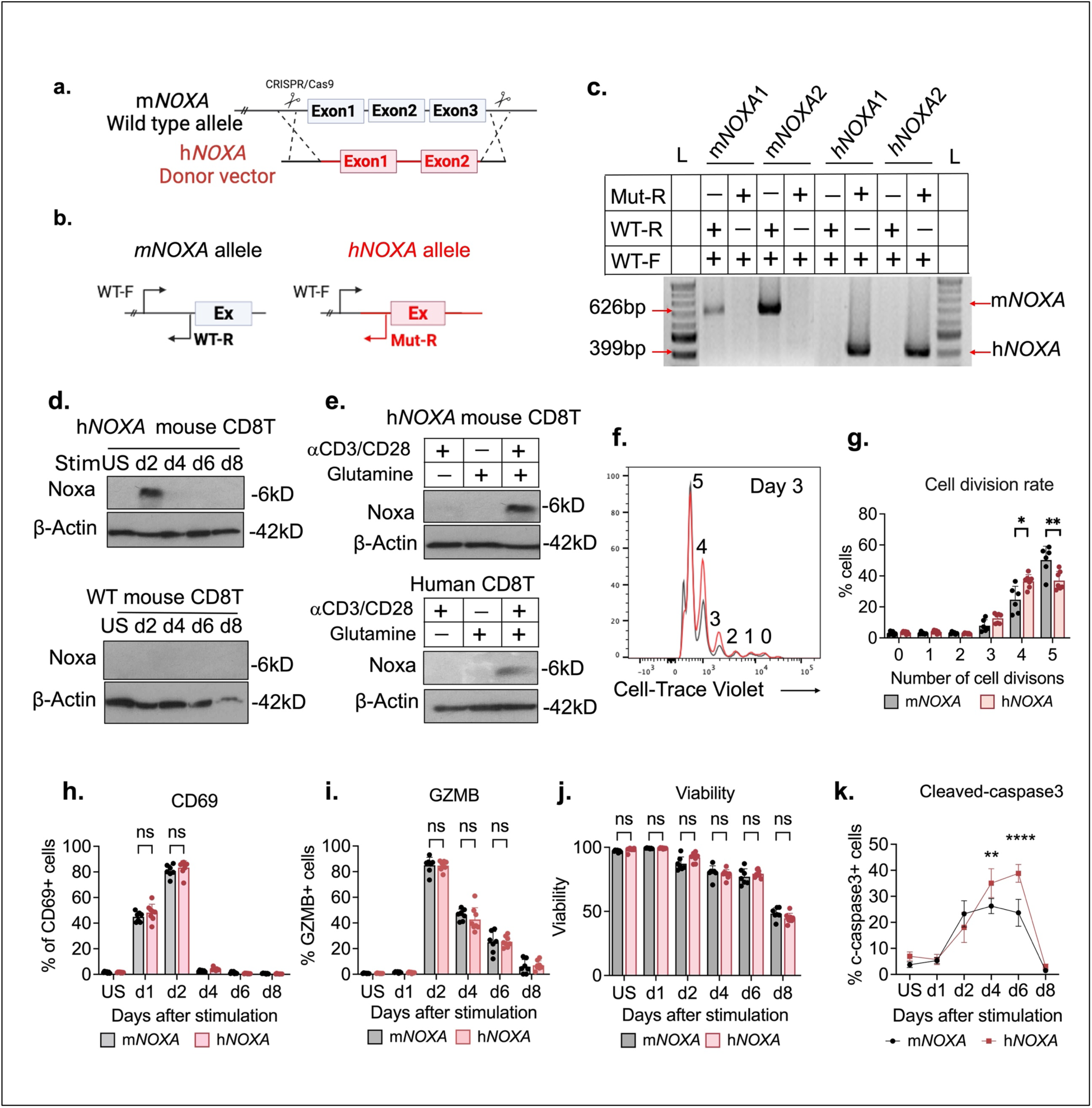
CD8^+^ T lymphocytes from gene replacement h*NOXA* mouse show few differences with wildtype T cells. **a.** Schematic representation of the replacement of the mouse *Pmaip1* (m*NOXA*) gene with the human *PMAIP1* (h*NOXA*) gene using CRISPR/Cas9-mediated genome editing. **b, c.** Genotyping of wild-type (m*NOXA*) and gene-replaced h*NOXA* homozygous mice by PCR using primers WT-F, WT-R, and Mut-R (**b**). PCR amplification of wild-type m*NOXA* alleles with WT-F and WT-R yields a 626 bp band, while amplification of h*NOXA* alleles with WT-F and Mut-R yields a 399 bp band (**c**). L – Quick-Load® 100 bp DNA Ladder (NEB). **d.** Western blot analysis of Noxa protein expression in CD8⁺ T cells isolated from the spleens of GR h*NOXA* and WT m*NOXA* mice at the indicated time points following in vitro stimulation with anti-CD3/CD28 beads. **e.** Noxa protein expression in h*NOXA* mouse CD8⁺ T cells (top panel) and naïve human CD8⁺ T cells (bottom panel) 48 hours after stimulation with anti-CD3/CD28 beads in high glutamine (HQ) or glutamine-free (NQ) medium. **f, g.** Representative overlaid histograms (**f**) of Cell-Trace Violet dilution on day 3 of stimulation of CD8⁺ T cells from m*NOXA* (grey) and h*NOXA* (red) mice, and quantification showing the frequency of cells in each doubling cycle (**g**) (n = 7). **h–k.** Percentage of CD69⁺ (**h**), Granzyme B⁺ (**i**) cells, viability (**j**), and cleaved-caspase-3⁺ cells (**k**) in isolated CD8⁺ T cells from m*NOXA* and h*NOXA* mice at the indicated time points following stimulation. (n=7)

Western blots of splenic CD8^+^ T cells from h*NOXA* or control m*NOXA* mice, stimulated with anti-mCD3/CD28 beads in vitro and probed with human Noxa-specific antibodies, showed that human Noxa was induced in h*NOXA* T cells as observed in human T cells, confirming the promoter was functioning normally, although induction was markedly transient compared to that observed in human CD8⁺ T cells stimulated in vitro (Fig. 6d, 1m, S1m). Additionally, induction of the human protein in CD8^+^ T cells from the GR mice was also glutamine dependent as in human CD8^+^ T cells (Fig. 6e, 3c), suggesting that early regulation of hNoxa was conserved in h*NOXA* T cells. Further characterization of the h*NOXA* and m*NOXA* CD8^+^ T cells that were stimulated in vitro revealed no significant differences in rates of activation or differentiation, and no differences in viability (Fig. 6h-j). However, h*NOXA* CD8^+^ T cells consistently proliferated at a slower rate than wildtype m*NOXA* CD8^+^ T cells and displayed higher levels of active (cleaved) caspase 3 at later times in their response to stimulation (Fig. 6f, g, k).

We also compared responses of GR h*NOXA* and control mice infected with lymphocytic choriomeningitis virus (LCMV). Analysis of spleen, liver, and small intestine intraepithelial lymphocytes (SI IELs) from the infected mice 12 weeks post-infection revealed no differences either in the percentage of CD8⁺ tetramers or in specific CD8⁺ T cell subsets in the spleen, including long-lived effector cells (LLECs), central memory T cells (Tcm), and effector memory T cells (Tem). Similarly, no differences were observed in non-lymphoid tissues, including SI IELs and liver (S6b-f).

Thus, these results suggest that Noxa is unlikely to be a driver in the response to either in vitro co-stimulation or in vivo acute infection in mouse CD8^+^ T cells. The consistently lower proliferation rate of the CD8^+^ T cells from the h*NOXA* mice, however, indicated possible growth regulation by the human protein, and merit further investigation.

### RNA-Seq analysis of activated CD8^+^ T cells from h*NOXA* mice reveals association with a proliferative gene signature

Studies in activated human T cells, described here (Fig. 3, 5) as well as previous published work in T leukemia cells^11, 13^, have established a role for Noxa in growth-related metabolic reprogramming. A growth-promoting effect on murine CD8^+^ T cells, specifically attributable to human Noxa (Fig. 6f, g), therefore, offered an opportunity to gather further insights into this non-canonical function which makes hNoxa a unique BH3-only protein. We were specifically interested in the gene expression profile of h*NOXA* T cells compared to those of control mice immediately following Noxa induction i.e., within the first two days of antigen encounter.

To identify pathways and targets responsive to hNoxa expression, we gathered RNA-Seq data from splenic h*NOXA* and control mouse CD8^+^ T cells co-stimulated in vitro with anti-CD3/CD28 for 48h. Transcriptomic profiling identified 118 significantly differentially expressed genes (DEGs), with 34 genes upregulated and 84 downregulated in h*NOXA* CD8⁺ T cells compared to their m*NOXA* counterparts (Fig. 7a). To focus on the impact of hNoxa in growth-related pathways, a hallmark gene set was preferred over available CD8^+^T-related gene sets for bioinformatics analysis. The virtual absence of representation of the murine *Pmaip1* (m*NOXA*) transcript in the h*NOXA* transcriptome in the volcano plot and heatmap of apoptosis-related transcripts provided additional validation of successful gene replacement (Fig 7a, e). Strikingly, gene set enrichment analyses (GSEA) of the data, and associated heatmaps, point to significantly reduced inflammation, hypoxia, ROS, and apoptosis-related gene signatures in the h*NOXA* CD8⁺ T cells (Fig 7b, c, e, S7a-c). TGF-β signaling, p53 and IL-2/STAT5 were also dominant in mouse CD8^+^ T cells at two days of stimulation compared to h*NOXA* CD8^+^T cells. Instead, OxPhos, G2/M checkpoint and cMyc-related proliferative gene signatures were enriched in the latter (Fig 7b, d, S7d). Given the largely similar responses of h*NOXA* and m*NOXA* CD8^+^ T cells to antigenic stimulation, the lower inflammation, hypoxia, ROS, and apoptosis-related signatures may reflect the reduced proportion of these transcripts in the total hNoxa transcriptome at 48 hours, rather than an actual reduction in transcript levels.

**Figure 7.**
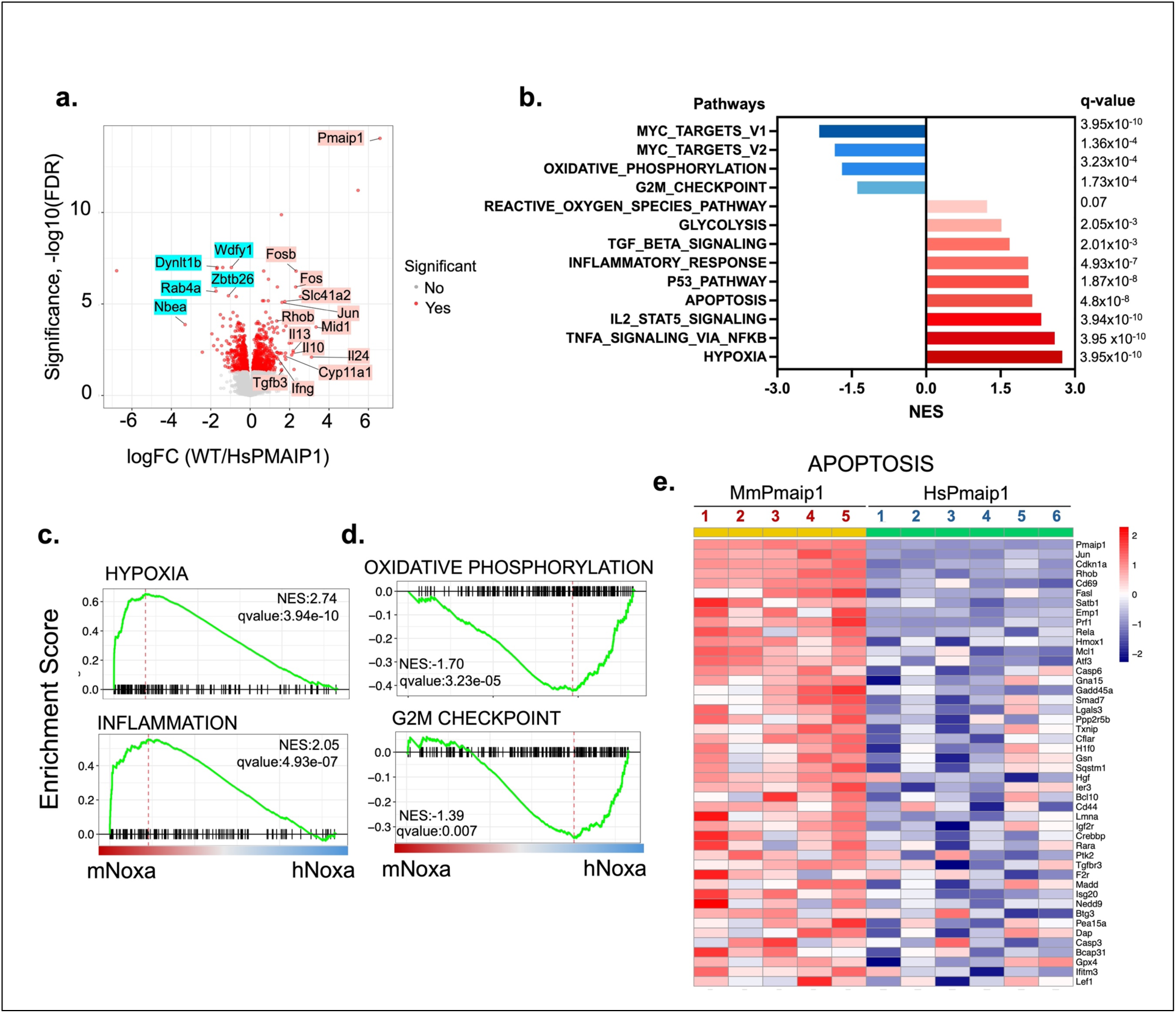
RNA-Seq analyses of activated CD8^+^ T cells from h*NOXA* mice reveal early association with a proliferative gene signature. **a.** Volcano plot displaying 118 differentially expressed genes between mNoxa and hNoxa expressing CD8⁺ T cells after 48 hours of stimulation. Selected genes overexpressed in 48h-stimulated CD8^+^ T are highlighted; mNoxa (pink) and hNoxa (seagreen). **b.** The most significantly altered pathways based on enrichment of differentially expressed genes (m*NOXA* (red) versus h*NOXA* (blue)) in GSEA Hallmark gene sets (FDR < 0.25). The y axis represents the enriched gene sets (either positive or negative), and the x axis represents the normalized enrichment scores (NES) for each gene set. **c**.**d.** Representative Gene Set Enrichment Analysis (GSEA) plots from the Hallmark gene set (hypoxia, inflammation, oxidative phosphorylation and G2M checkpoint) of RNA-seq data from 48h stimulated m*NOXA* and h*NOXA* CD8^+^ T cells (m*NOXA* versus h*NOXA*). Normalized enrichment score (NES) and nominal p value are indicated. **e.** Heatmap of the top significantly differentially expressed genes (DEGs) (n = 46) in the apoptosis pathway comparing 48-hour-stimulated m*NOXA* (MmPmaip1) with h*NOXA* (HsPmaip1) CD8⁺ T cells, where n indicates the number of leading-edge genes.

Overall, these results support a model in which human Noxa promotes a growth and proliferative phenotype in activated CD8⁺ T cells upon its induction. Interestingly, some of the upregulated transcripts in murine h*NOXA* CD8^+^ T cells code for proteins associated with mitochondrial metabolism, and others share conserved domains that are involved in protein-protein interactions, intracellular vesicle trafficking and localization (Fig 7a). Implications of the transcriptomic profiling results for hNoxa function will be further discussed in the following section.

## DISCUSSION

We have investigated the role of human BH3-only Bcl-2 protein, Noxa, in the CD8^+^ T cell response to antigenic stimulation using an in vitro model of TCR co-stimulation. Human Noxa, like all BH3-only proteins, bears little resemblance to multidomain members of the Bcl-2 family beyond the shared BH3 domain^5^. However, it also stands apart from its pro-apoptotic BH3-only counterparts and its own murine homologue in its ability to regulate both cell growth as well as death in immune cells. Our study reveals that, in human CD8^+^ T cells responding to antigenic stimulation, Noxa contributes to the metabolic reprogramming that precedes expansion as well as to the apoptotic death that terminates the response without affecting their differentiation or cytotoxic potential.

The in vitro model we have described captures the entire course of a human CD8⁺ T-cell immune response under a single antigen exposure, without withdrawal or repeated stimulation, unlike short-term human T cell stimulations or exhaustion models. While the model lacks the complexity of the intact immune system and focuses on a single T-cell subtype, it has allowed us to track, in real time, activation, proliferation, differentiation and apoptotic phases as well as memory formation in freshly isolated CD8^+^ T cell populations in response to antigen, and to correlate Noxa expression with the phases of the response (Fig 1n). The differences between the human and murine versions of Noxa were the primary impetus for developing an in vitro co-stimulation assay to determine hNoxa’s role in T cells. A commercial antibody suitable for detection of hNoxa by flow cytometry is currently unavailable, however, the high yields of naïve cells from a single human donor as well as the superior proliferative capacity of stimulated CD8^+^ T cells in vitro made detection of Noxa in cell extracts by western blot a feasible approach. The absence of detectable Noxa in naïve and memory CD8^+^ T populations and its rapid transcriptional induction following anti-CD3/CD28 co-stimulation prior to their entry into the expansion phase pointed to tight regulation and a coordinated involvement of the protein with early and late phases of the CD8^+^ T immune response (Fig 1m, n).

The in vitro stimulation experiment consistently produced a population of cells with distinct memory-like characteristics, while revealing the limitations of relying on conventional memory T-cell markers such as CD45RO/CD45RA, which are primarily reporters of antigen encounter rather than of stable commitment to the memory lineage^21, 22^. Responding CD8⁺ T cells displayed a stable CD45RO⁺/CD45RA⁻ phenotype while still maintaining high effector activity; however, a sudden, significant drop in levels of the late activation marker, HLA-DR^23, 24^ at the end of the immune response coincided with the return of activation, effector, and cytotoxic markers to baseline, signaling a shift away from effector activity and toward memory formation. The viable cells remaining at this stage display several distinguishing characteristics of memory cells^4, 43^, including increased mitochondrial content and membrane potential, robust spare respiratory capacity (indicative of mitochondrial fitness), a dependence on FAO for mitochondrial activity in the quiescent state, and an accelerated response to recall stimulation (Fig 2). We conclude that declining HLA-DR expression is a more reliable marker for the effector-to-memory transition than CD45RO/CD45RA ratios in a short-term in vitro assay. The long-term functionality and self-renewal capacity of these memory-like cells remain to be explored but taken together, the current data establish our system as a tractable in vitro platform for dissecting human CD8^+^ T cell response to antigenic stimulation and for refining strategies to produce functionally competent, memory-biased T cells.

Transcription of the *NOXA* gene is activated in CD8⁺ T cells in response to stimulation, but protein translation requires the presence of glutamine (Fig 3). This requirement for glutamine is independent of canonical glutaminolysis as Noxa induction was unaffected by chemical inhibition of glutaminase, and supplementing glutamine-free medium with glutamate or αKG, breakdown products of glutaminolysis, did not restore Noxa protein expression (Fig 3f, k, 5a). In fact, our data showed that glutamine-dependent translation of *NOXA* mRNA was a direct consequence of glutamine regulation of mTORC1 activity (Fig 4). This study is the first demonstration of the involvement of mTORC1 in regulating the metabolic switch to glutaminolysis in TCR activated human CD8^+^ T cells.

The effects of short-term inhibition of mTOR in impairing mitochondrial function contrast with the effects of sustained inhibition of mTORC1, which promote FAO and memory phenotypes with a higher SRC^40, 44^. The differential effects can be explained in part by mTORC1’s role in regulating the key metabolic transcription factor, cMyc, essential for glycolysis and glutamine metabolism^45, 46^. Acute short-term inhibition of mTORC1 likely affects both glycolytic and glutamine-driven metabolic pathways, impairing mitochondrial fitness and bioenergetic capacity at a critical stage of T cell priming. Thus, the negative effects of glutamine loss or deprivation on both Noxa expression and mitochondrial respiration (Fig 3g) were directly attributable to inhibition of mTORC1 activity. This dependence on mTOR activity for expression contrasts with that of Bim, a BH3-only Bcl-2 family member that is involved in central and peripheral deletion of T lymphocytes and induced in the presence of rapamycin^47^.

Glutamine deficiency compromised mitochondrial SRC and prevented Noxa protein expression in stimulated CD8^+^ T cells. Despite this, loss of Noxa in glutamine-replete medium through RNA interference or CRISPR editing enhanced, rather than impaired, mitochondrial respiratory activity. Additionally, intracellular glutamate levels were significantly reduced in the stimulated *NOXA* knockdown and knockout populations. Given the similar outcomes in the CD8^+^ T cell response to chemical inhibition of GLS, these data point to a role for Noxa in regulating GLS activity (Fig 5a-e, g-i). The enhanced respiratory activity - despite the reduction in glutamate levels following Noxa loss - suggests that cells likely switch to an alternate, anaplerotic, carbon source^42^ to replenish TCA cycle intermediates and generate energy for growth and expansion. One possible route is through increased activity of pyruvate carboxylase, a major anaplerotic enzyme that directly generates oxaloacetate from pyruvate in the mitochondria. These observations parallel murine studies showing GLS deletion did not impair mitochondrial function or effector activity, even during chronic infection, and stimulated GLS deficient T cells demonstrated metabolic flexibility in rerouting substrates for proliferation and differentiation^48^. A related study points to a role for glutamine restriction during primary activation of murine CD8⁺ T cells in enhancing glycolysis and SRC, and in preventing premature exhaustion and improving persistence without loss of anti-tumor effector function^49^. Similarly, *NOXA*-deficient human CD8⁺ T cells also displayed increased viability and greater persistence compared to control cells, while remaining functionally effective (Fig 5). It is intriguing that Noxa is needed for directing glutamine to the TCA cycle but knocking it out enhances, rather than reduces, mitochondrial fitness and the cells’ ability to grow.

A major challenge in investigating the unique dual functions of hNoxa had been the absence of an in vivo model to validate in vitro findings, given the multiple differences between the murine and human Noxa proteins. The gene replacement model described here is the first to enable the study of hNoxa in an in vivo context. Initial characterization showed that the responses to both in vivo infection and in vitro TCR co-stimulation were similar in h*NOXA* and wildtype murine CD8^+^ T cells, suggesting that the Noxa protein was unlikely to be a driver of this response, although caspase-3 activity was higher and peaked later in GR h*NOXA* CD8^+^ T cells in the in vitro assay (Fig 6 and S6). It should be noted that the average duration of a human CD8^+^ T cell in vitro response to TCR co-stimulation, as described in our study, is 19 days which includes a longer expansion phase and later onset of apoptosis compared to an equivalent mouse CD8^+^ T in vitro assay which lasted 8 days. An antibody against mNoxa protein is currently unavailable but induction of the m*NOXA* transcript typically occurs on day 3 of the 8-day response^50^ compared to 24-40h timeline for the human transcript and protein, which persists for the duration of the response (Fig 1), underscoring the growth promoting role of the latter.

Initial characterization of CD8^+^ T cells from the GR model reaffirmed hNoxa’s dependence on glutamine and early association with a proliferative rather than cytotoxic or inflammatory gene expression signature following antigen exposure (Fig 6ab, 7). The enriched G2/M signature (Fig 7d) highlighting genes at the G2/M checkpoint where cells check for DNA damage before entering mitosis, provides additional support for a proliferative role through cell-cycle control, and may explain the observed increase in doubling times of hNoxa cells (Fig. 6f, g). Several significantly upregulated transcripts in h*NOXA* T cells encode proteins with conserved domains, such as zinc finger, FYVE, PH, and WD40 domains, largely associated with signaling via protein-protein interactions, protein transport, membrane localization or vesicle trafficking (Nbea, Rab4a, Dynlt1b, and Wdfy1)^51–54^ with the exception of Zbtb26, a zinc finger and BTB domain-containing member of the Zbtb family of transcription factors known to regulate T cell differentiation^55^. Further studies will be required to determine whether human Noxa directly regulates transcriptional programs or modulates these domain interactions indirectly, and how they would contribute to hNoxa function. Future studies could also include similar RNA-Seq analyses to compare expression patterns of stimulated wildtype with matched donor *NOXA* KO human CD8^+^ T cells.

We have established a comprehensive in vitro immune response platform in human CD8^+^ T cells, which has provided insights into Noxa’s divergent roles in early and late phases that would have been challenging to identify in an in vivo study. In T leukemia cells the switch between the pro-survival/metabolic and pro-apoptotic functions of hNoxa is regulated by phosphorylation of S^13^, which inhibits its pro-apoptotic activity while promoting its role as a metabolic regulator of growth, by preventing its interaction with Mcl-1^11–13^. Further studies are needed to determine whether phosphoregulation governs the transition of Noxa from a metabolic regulator to a death promoter in a similar manner in primary T cells. It is notable that the growth promoting metabolic function of hNoxa has only been identified in cells of the immune system; Noxa’s role in epithelial cells is primarily that of a pro-apoptotic BH3-only protein^14, 56^.

Our findings offer strong rationale for engineering *NOXA* to improve efficacy and durability of adoptive immunotherapies. Noxa is unique in enabling both growth-promoting metabolic reprogramming and growth-suppressing apoptosis in CD8^+^T cells in response to immune challenge and knocking out the *NOXA* gene in stimulated CD8+T cells delays the apoptotic death that occurs in the T cells at the end of an immune response, without compromising their cytotoxic potential. While engineered knockout of other pro-apoptotic BH3-only Bcl-2 genes, such as *BIM*, can also be predicted to extend cell viability, other BH3-only proteins lack Noxa’s ability to regulate glutamine metabolism. One study shows association between Bim and glutamine in multiple myeloma where glutamine withdrawal, in fact, increases Bim expression and promotes apoptotic interactions with Bcl-2^57^. Our studies suggest that targeting *NOXA* could enhance the durability and persistence of CAR-T and TIL therapies, particularly within nutrient-restricted tumor microenvironments where metabolic stress and apoptosis limit their efficacy.

## METHODS

### Isolation, culture and expansion of human CD8+ T cells from normal donors

Fresh, de-identified human, donor peripheral blood mononuclear cells (PBMCs) recovered from leukoreduction system chambers (TRIMA Cones), obtained from Memorial Blood Center (St. Paul, MN) were subjected to red blood cell lysis using RBC lysis buffer (Biolegend, 420302) per manufacturer’s instructions. ImmunoCult™-XF T Cell Expansion Medium (STEMCELL Technologies Cat# 10981), Naïve or total CD8⁺ T cell populations were isolated using the EasySep™ Human CD8⁺ T Cell Isolation Kit (STEMCELL Technologies Cat# 17953 and Cat# 17968) according to the manufacturer’s protocol. Purified CD8⁺ T cells were cultured at a density of 1 × 10⁶ cells/mL in ImmunoCult™-XF medium and stimulated with ImmunoCult™ Human CD3/CD28 T Cell Activator (STEMCELL Technologies Cat# 10971) 4 ng/mL human IL-2 (66 IU/ng) (Peprotech, AF-200-02-100). Unless otherwise indicated, cells were maintained at 37 °C in a humidified incubator with 5% CO₂, and fresh medium was added as needed to maintain cell density and viability.

### Generation of human CD8+ memory T cells and re-stimulation

CD8^+^ T cells, activated as described above were maintained without further stimulation, and with fresh media supplementation, as needed. Viable cells, T cells, were isolated by density gradient centrifugation using Ficoll-Paque™ PREMIUM density gradient media (Cytiva, 17544602) 17–19 days post-activation when of the cell population recorded a significant drop in HLA-DR mean fluorescence intensity (MFI). Recovered cells, defined as the memory-like population, were allowed to rest for 2 hours in ImmunoCult™-XF T Cell Expansion Medium, Memory-like T cells, like CD8⁺ T cells, were re-stimulated as described above for naïve and total CD8T cells and evaluated for Noxa for activation, expansion, differentiation and metabolic phenotype.

### Real-Time Quantitative PCR

For *Noxa* gene expression analysis, total RNA was extracted from human CD8⁺ T cells and reverse-transcribed into cDNA using the Cells-to-cDNA™ II Kit (Thermofisher Scientific, AM1722). Quantitative real-time PCR (RT–qPCR) was performed using PowerTrack SYBR Green Master Mix (Thermo Fisher Scientific, A46012) on a QuantStudio 3 Real-Time PCR System (Thermo Fisher Scientific). Expression levels were normalized to the human housekeeping gene *ACTB* (β-actin), and relative expression was calculated using the ΔΔCt method. Primer sequences were as follows: human Noxa (PMAIP1), forward 5′-ATGCCTGGGGAAGAAGCGCGCAAG-3′ and reverse 5′-GGTTCCTGAGCAGAAGAGTTTGGA-3′; ACTB, forward 5′-TCCCTGGAGAAGAGCTACGA-3′ and reverse 5′-AGCACTGTGTTGGCGTACAG-3′.

### Inhibitor treatment

For studies on the regulation of glutamine metabolism and mTOR signaling, CD8⁺ T cells were treated with inhibitors at the time of activation. Cells were cultured in T Cell Expansion Basal Medium (TCEM; Gibco, A1048501) supplemented with OpTmizer CTS and CTS Immune Cell Serum Replacement (Gibco, A2596101), in which L-glutamine was adjusted to high (2 mM), low (0.2 mM) or glutamine-free (0 mM) conditions. The following compounds were used at the indicated final concentrations: CB-839 (2 µM; MedChemExpress, HY-12248), 6-diazo-5-oxo-L-norleucine (DON; 200 µM; Sigma-Aldrich, D2141), rapamycin (100 nM; Cayman Chemical, 13346), everolimus (100 nM; Cayman Chemical, 11597), V-9302 (2 µM; ASCT2 inhibitor; Cayman Chemical, 27688), JPH203 (1 µM; LAT1 inhibitor; Cayman Chemical, 29715) and N-ethylmaleimide (NEM; 100 µM; Cayman Chemical, 19938). All inhibitors were prepared as stock solutions and diluted in culture medium immediately before use; vehicle-treated cells served as controls. To assess the contribution of fatty acid oxidation in memory-like CD8⁺ T cells, cells were treated with etomoxir (5 µM; CPT1a inhibitor; Sigma-Aldrich, 5094550001) at the time of re-stimulation.

### Western blotting

Cells were washed once with ice-cold PBS and lysed in RIPA buffer supplemented with protease and phosphatase inhibitor cocktails (Thermo Fisher Scientific, 89900). Lysates were clarified by centrifugation at 14,000 × g for 15 min at 4 °C, and protein concentration was determined using a BCA protein assay kit (Thermo Fisher Scientific, 23225), according to the manufacturer’s instructions. Equal amounts of protein (40 µg) were mixed with 2× Laemmli sample buffer (Bio-Rad, 1610737), heated at 95 °C for 3 min, resolved by SDS–PAGE on 8–16% Tris–glycine gels (Bio-Rad, 5671104) and transferred to nitrocellulose membranes (Cytiva, 10600009). Membranes were blocked in 5% milk in TBS-T (20 mM Tris-HCl, pH 7.6, 137 mM NaCl, 0.1% Tween-20) and incubated overnight at 4°C with primary antibodies diluted in TBS-T buffer. The following primary antibodies were used: anti-β-Actin (Santa Cruz, sc-69879, 1:7000), anti-Bim (Cell Signaling, 2933, 1:2000), anti-Mcl-1 (Cell Signaling, 94296, 1:1000), and anti-Noxa (Santa Cruz, sc-56169, 1:250; Cell Signaling, 14766, 1:1000). After washing, membranes were incubated with appropriate HRP-conjugated secondary antibodies (Genesee Scientific, 20-303 and 20-304) for 1 hour at room temperature. Signal was developed using Pierce™ ECL Western Blotting Substrate (Thermofisher Scientific, PI80196) and films (Cytiva, 28906839; Genesee Scientific, 30-810). For reprobing, membranes were stripped in low pH TBS-T buffer (pH 2.5–3.0) for 5 minutes, followed by re-blocking before re-incubation with antibodies, as described previously^11^.

### Human CD8^+^ T cell flow cytometry (surface and intracellular staining)

Single-cell suspensions of 5 × 10⁵ human CD8⁺ T cells were collected at indicated time points post-stimulation for flow cytometric analysis. Viability was assessed using Live/Dead Fixable Near-IR Dead Cell Stain (Biolegend, 423106), followed by surface marker staining: CD8 (Biolegend, 344740 or 344776), CD69 (Biolegend, 310938), CD62L (Biolegend, 304832), CD107a (Biolegend, 328626), HLA-DR (Biolegend, 327006), CD45RO (Biolegend, 304206) and CD45RA (Biolegend, 304156) for 30 min at 25 °C in the dark. For intracellular staining, cells were fixed and permeabilized using the FoxP3/Transcription Factor Fixation/Permeabilization Buffer Set (Cell Signaling, 43481) and subsequently incubated with intracellular antibodies: Granzyme B (Biolegend, 515406), Cleaved Caspase-3 (Cell Signaling, 8788), Phospho-4E-BP1 (Cell Signaling, 7547S) and Phospho-S6 Ribosomal Protein (Cell Signaling, 5044S) for 45 min at 25 °C in the dark. All surface and intracellular antibodies were used at 1:100. Samples and single stain controls were washed in FACS buffer (2% FBS, 0.1% Sodium Azide in 1x PBS). For proliferation assays, cells were labeled with CellTrace™ Violet (Thermofisher Scientific, C34571). Compensation beads (Invitrogen, 01-2222-42 and 01-3333-42) were used to generate single-color controls for compensation. Data acquisition was performed on a BD LSR II or BD FACSymphony™ A3 flow cytometer, and data were analyzed using FlowJo software (version 10.8.1; BD Biosciences). Gating proceeded as follows: (1) lymphocytes by FSC-A vs SSC-A; (2) live cell subsets were selected by ZombieNIR^+^ exclusion; (3) CD8^+^ T-cell subsets were gated off ZombieNIR - subsets; (4) functional/phenotypic markers (e.g., CD69, CD62L, CD107a, HLA-DR, GZMB, cleaved-caspase3) quantified within the viable CD8^+^ gate. All isolated naïve CD8+ T cells (CD8^+^CD45RA^+^CCR7^+^) or total CD8+ T cells (CD8^+^) cells displayed >95% purity prior to downstream activation analysis and subsequent experiments. Compensation was performed using single-stained controls, and gating thresholds were defined using fluorescence minus one (FMO) controls or based on marker expression levels in unstimulated CD8^+^ T cells. A minimum of 1 × 10^4^ CD8^+^ events per sample were acquired on a digital flow cytometer.

### Seahorse Live Cell Metabolic Flux Assays

The analysis of Extracellular Acidification Rate (ECAR) and Oxygen Consumption Rate (OCR) of CD8+ T cells was performed using the Seahorse XFe96 Analyzer (Agilent). Seahorse XF96 cell culture microplates (Agilent) were coated with Cell-Tak (Corning, 354240) according to the manufacturer’s instructions. Human CD8⁺ T cells were resuspended in Seahorse XF Base RPMI Medium (Agilent, 103576-100) supplemented with 10 mM glucose, 2 mM glutamine, and 2 mM sodium pyruvate (all from Agilent), adjusted to pH 7.4. A total of 2 × 10⁵ cells per well were plated, centrifuged to promote adherence, and incubated at 37°C in a non-CO₂ incubator for 1hr prior to analysis. The perturbation profiling of metabolic pathways was achieved with Agilent Seahorse XF Mito Stress Test (Agilent, 103015-100): 1.5 µM oligomycin, 2.0 µM FCCP, and 0.5 µM rotenone/antimycin A. Brightfield imaging prior to assay initiation and post-assay Hoechst 33342 staining (1:500) allowed for data normalization using a Cytation 1 plate reader (BioTek). Spare respiratory capacity was expressed as the difference between maximal OCR and basal OCR.

### siRNA transfection

Validated human Noxa-targeting siRNA and non-targeting control siRNAs were purchased from Horizon Biosciences (Human PMAIP1 siRNA, L-005275-00-0005; Non-targeting Control siRNA, D-001810-10-05). Unstimulated human CD8⁺ T cells (5 × 10⁶ cells) were resuspended in 200 µL T buffer (Invitrogen) and electroporated with 12 µL siRNA (100 µM) using the Neon Transfection System (Invitrogen) at 2150 V, 20 ms pulse width, and 1 pulse, as described previously^58^. Following electroporation, cells were cultured in RPMI 1640 (HyClone, SH30027.02) supplemented with 10% FBS (Atlas Biologicals, FS-0050-AD) and 2 mM L-glutamine (Thermofisher Scientific, 25030081). Four hours post-transfection, CD8⁺ T cells were activated by plate-bound stimulation using anti-CD3 and anti-CD28 antibodies (Biolegend, 317325 and 302934). 1% Penicillin-Streptomycin (Gibco, 15140122) was supplemented 48hr after stimulation.

### CRISPR/Cas9 Genomic Editing

Human CD8+T cells (2x10^6^) were activated and incubated in ImmunoCult™-XF T Cell Expansion Medium for 48h prior to transfection with sgRNAs. Briefly, 100pmol sgRNAs (Synthego, Noxa-targeting sgRNA: CACCGGCGGAGAUGCCU) were incubated with 1ug of CleanCap^R^ Cas9 mRNA (TriLink BioTechnologies, L-7606) for 10-15 min at room temperature. Preactivated CD8+ T cells were washed once in cold PBS and resuspended in 19.5 ul T buffer (Invitrogen). Neon Transfection System (Invitrogen) was used for the transfection experiments, and each set of experiments was conducted via the 10ul tips (Invitrogen). 1400 Volts, 10 pulse widths, 3 pulses were the setting for the parameters, as described previously^59^. After transfection, the cells were transferred to 1ml T cell expansion basal medium (TCEM) (Gibco, A1048501), with Optimizer CTS, CTS Immune Cell SR (Gibco, A2596101) and 2mM L-Glutamine. 1% Penicillin-Streptomycin (Gibco, 15140122) were added 48h after transfection.

### Glutamate assay

Glutamate levels were quantified using the fluorometric Glutamate Assay Kit (Abcam, ab138883) following the manufacturer’s instructions. Briefly, CD8⁺ T cells were stimulated and treated with indicated concentrations of CB-839 or vehicle control for 48 h. After treatment, cells were harvested, counted for normalization, and lysed in RIPA buffer supplemented with protease and phosphatase inhibitors. Lysates were cleared by centrifugation at 14,000 × g for 15 min at 4°C. Protein concentrations were determined using the Pierce™ BCA Protein Assay Kit (ThermoFisher Scientific). 10ug of total protein per sample were diluted to 50 µL with assay dilution buffer and loaded into black 96-well plates, followed by the addition of 50 µL of reaction mix. The reaction was incubated at room temperature, and fluorescence (Ex/Em = 540/590 nm) was measured at multiple time points (30, 45, 60, 75, and 90 min) using an Infinite M1000 PRO Microplate Reader (Tecan).

### IL-2 ELISA

IL-2 secretion was quantified using the ELISA MAX Deluxe Set Human IL-2 (BioLegend, 431804) according to the manufacturer’s instructions. Briefly, CD8⁺ T cells were stimulated as indicated, and culture supernatants were collected at the specified time points from wells containing 1 × 10⁶ cells. Supernatants were clarified by brief centrifugation and stored at −80 °C until analysis. Standards and samples were added in duplicate to coated plates and incubated according to the kit protocol, followed by detection antibody, streptavidin–HRP and substrate development. Absorbance was measured at 450 nm with a reference wavelength of 570 nm on a Cytation 1 plate reader (BioTek), and IL-2 concentrations were calculated from a standard curve and normalized to the number of cells per well.

### Animal studies

All animal procedures were performed under protocols approved by the Institutional Animal Care and Use Committee (IACUC) at the University of Minnesota. Human Noxa gene-replacement C57BL/6 mice (hNoxa knock-in) were generated by Beijing Biocytogen Co., Ltd. Wild-type C57BL/6J mice (Jackson Laboratory, stock no. 000664), 8–12 weeks of age, male and female, were used as controls. For genotyping of *hNoxa* gene-replacement mice, genomic DNA was isolated from ear or tail biopsies using the DNeasy Blood & Tissue Kit (QIAGEN, 69504). PCR was performed using gene-specific primers to distinguish wild-type and knock-in alleles: Genotyping primers for mNoxa: Forward: GCTCCTCCGAATTTACCACACT, Reverse: AACCTGGCTCTGGGAAGTTTGGTCG; Genotyping primers for hNoxa: Forward: GCTCCTCCGAATTTACCACACT, Reverse: CGCAGGAAGCACACTGGAGATGCTG. Mice were housed under specific pathogen-free conditions in microisolator cages with ad libitum access to autoclaved food and water. LCMV Armstrong infections were performed by intraperitoneal (i.p.) injection of 2 × 10⁵ PFU in a BSL-2 animal facility.

### Purification and in vitro stimulation of mouse CD8+ T cells

Single-cell suspensions were prepared from spleens of 4- to 6-week-old male and female hNoxa or mNoxa mice by mechanical dissociation through 70-μm nylon cell strainers (BD Biosciences), followed by red blood cell lysis using 1× RBC Lysis Buffer (BioLegend, 420302). Naive CD8+ T cells were isolated using the Mouse CD8+ T Cell Isolation Kit (Miltenyi Biotec, 130-104-075), according to the manufacturer’s instructions. Purified CD8+ T cells were cultured in T Cell Expansion Basal Medium (TCEM) supplemented with CTS™ Immune Cell Serum Replacement (Thermo Fisher Scientific), 2 mM L-glutamine, 100 μg/mL streptomycin, 100 U/mL penicillin, and 100 IU/mL recombinant human IL-2. For in vitro activation, cells were stimulated with Dynabeads™ Mouse T-Activator CD3/CD28 (Thermofisher Scientific, 11452D) at a bead-to-cell ratio of 1:1, unless otherwise indicated.

### LCMV infection and tissue harvest

Mice were infected with 2 × 10⁵ PFU of LCMV Armstrong strain via intraperitoneal injection in a BSL-2 animal facility. At 12 weeks post-infection, mice were euthanized following retro-orbital injection of 3 μg anti-CD8α antibody (53–6.7) (Cytek Biosciences, SKU 35-0081-U025 and SKU 35-0081-U025) administered 5 minutes prior to sacrifice.

Peripheral blood was collected by cheek bleed into heparinized tubes, and mice were sacrificed for tissue harvest. Inguinal lymph nodes (LNs) and spleens were collected into RPMI 1640 supplemented with 5% FBS and passed through 70-µm cell strainers.

For intraepithelial lymphocyte (IEL) isolation, small intestines were excised, cleared of fat, flushed of luminal contents, and immersed in IEL buffer (1× HBSS supplemented with Hepes, sodium bicarbonate, and 2% FBS). Peyer’s patches were removed, intestines were opened longitudinally, vortexed, and kept on ice in 20 mL IEL buffer. After decanting the supernatant, tissue was washed with 30 mL fresh IEL buffer, transferred to 50 mL Erlenmeyer flasks containing 30 mL IEL buffer supplemented with 5% FBS and 154 g/L dithioerythritol (DTE) (Millipore, 233152), and stirred at 37°C for 30 min on a Variomag Poly 15 (Thermo Fisher Scientific). The supernatant was filtered through a 70-µm strainer. This extraction was repeated once, and supernatants were pooled.

For liver lymphocyte isolation, livers were excised (excluding the gallbladder), placed in 5 mL harvest medium on ice, dissociated using GentleMACS C tubes (Miltenyi Biotec) on a GentleMACS Dissociator (program: m_spleen_01.01, twice), and filtered through 70-µm strainers. Non-lymphoid tissue (NLT) cell suspensions were pelleted and resuspended in 5 mL of 44% Percoll (diluted in RPMI 1640), overlaid with 3 mL of 67% Percoll (diluted in PBS), and centrifuged for 20 min at 800 × g at room temperature with minimal acceleration and deceleration. Mononuclear cells were collected from the interface, washed, and used for downstream analyses. Red blood cells in blood and spleen samples were lysed using ACK lysis buffer (150 mM ammonium chloride, 1 mM potassium bicarbonate, 0.1 mM EDTA in water) prior to staining.

### Tetramer reagents and mouse CD8+ T cell flow cytometry

Tetramers were made using biotinylated monomers (H-2D^b^ KAVYNFATM [gp33/D^b^]; H-2D^b^ SGVENPGGYCL [gp276/D^b^]; H-2D^b^ FQPQNGQFI [NP396/D^b^]) from the NIH Tetramer Core at Emory University. For gp33, gp276. and NP396, R-PE-streptavidin (ThermoFisher Scientific, S21388) was used. Fluorophore-conjugated streptavidin was added to 20 μg of monomer, in 10 additions of 3.18 μg, each 10 min after the other (at room temperature). Class II tetramer (APC-conjugated 2W:I-A^b^ EAWGALANWAVDSA tetramer) was a kind given from Dr. T Dileepan (University of Minnesota). Tetramer was stored at 4 °C before use.

Samples and single stain controls were washed in FACS buffer (2% FBS, 2 mM EDTA in 1x PBS), followed by Fc blocking (BD Pharmingen, 553142) for 5 min (1/400). Antibodies/viability dye for staining were then added for 60 min at RT with concurrent tetramer staining when used: CD4 (BD Horizon, 565974), CD8a (BD Horizon, 750024), TCR β Chain (BD Horizon, 562840), CD103 (BD Horizon, Cat# 563087), CD127 (BD Horizon, 612841), CD27 (BD Horizon, 563605), CD25 (BD Horizon,564021), KLRG1 (BD Horizon, 564014), CD62L (BD Horizon, 564109), CD44 (IM7) (Cytek Biosciences, SKU 80-0441-U025), CD185 (CXCR5) (Biolegend, 145511), CD183 (CXCR3) (Biolegend, 126513) and Ghost Dye™ Red 780 (Cytek Biosciences, SKU 13-0865-T500), CD69 (H1.2F3) (eBioscience, 17-0691-82).

All surface antibodies were used at 1/200, except anti-CD69 and anti-CD127 (1/100), and MHC class II tetramer (10nM). Viability dye was used at 1/1,000. Samples were acquired using Cytek Biosciences Aurora instrument. Singlet lymphocytes were gated by forward scatter area (FSC-A)/side scatter area (SSC-A), FSC-A/FSC-height (FSC-H), and SSC-A/SSC-height (SSC-H). Live, TCRβ+ cells were then gated according to CD4 and CD8α staining, and, in the latter, by I.V. labeling status (blood, I.V.-positive; LN, IEL, I.V.-negative; spleen, I.V.-low; liver, not gated by I.V. labeling status). Subsequently, cells were identified by tetramer binding (tet+ CD44hi). For subsetting of memory, blood, spleen, and LN, LCMV-specific (i.e. tetramer-binding) T cells were gated as KLRG1+/CD62L-(long-lived effector cells), KLRG1-/CD62L-(T effector memory), and KLRG1-/CD62L+ (T central memory). Resident cells in liver and IEL were identified as CD69+ or CD103+.

In vitro stimulate splenic mouse CD8+ T cells were tested following stimulation for flow cytometric analysis. Viability was assessed using Live/Dead Fixable Near-IR Dead Cell Stain (Biolegend, 423106), followed by surface marker staining: CD8a (Biolegend, 100711), CD69 (Biolegend, 104559), CD62L (Biolegend, 104417), CD44 (Biolegend, 103021), CD127 (Biolegend, 135021) and KLRG1(Biolegend, 138423); and with intracellular antibodies: Granzyme B (Biolegend, 515406), Cleaved Caspase-3 (Cell Signaling, 8788). All surface and intracellular antibodies were used at 1:100.

### Bioinformatics analysis of RNA sequencing data

Read mapping and quantification were performed using a custom MSI pipeline (v.1.0.1)^60^. This pipeline aligned fastq files to the GRCm39 reference genome using HiSAT2 (v.2.1.0) and quantified the aligned reads to the version 110 GRCm39 gtf from Ensembl^61^ using featureCounts (v.2.0.6)^62^. Differential expression testing and pathway analysis were performed in R (v.4.4.1)^63^. The quasi-likelihood test method was used for differential expression with the EdgeR package (v.4.2.1)^64^. The clusterProfiler package (v.4.12.6)^65^ was used for gene set enrichment analysis with DE lists ranked by p-value (-log10 with fold-change direction assigned). Volcano plots of differential gene expression and heatmaps of the leading GSEA leading edge genes were created using the R ggplot2 package (v.3.5.1)^66^ and pheatmap package (v.1.0.12)^67^, respectively.

### Quantification and statistical analysis

All experiments were repeated independently at least three times with similar results. The number of samples (n) were described in detail for each figure panel. Data are presented as mean ± standard error of the mean (SEM) for biological replicates unless otherwise specified. Statistical analyses were performed using two tailed Student’s t-test for comparison between two independent groups and one-way or two-way analysis of variance (ANOVA) for multiple comparisons using GraphPad Prism 10. In all figures, statistical significance is marked as follows: not significant (ns), ∗P < 0.05, ∗∗P < 0.01, ∗∗∗P < 0.001 and ∗∗∗∗P < 0.0001.

## Author contributions

T.Y. and A.K. conceptualized and designed the study and wrote the manuscript. T.Y. performed most of the experiments and data analyses. H.S. contributed to the glutamine and mTOR studies and edited the manuscript. J.M.B. performed the pan T cell activation experiment. W.J.V. performed and analyzed the GR mouse infection studies. R.L. performed the bioinformatics analysis of the RNA-Seq data and wrote the corresponding methods section. B.W. and B.M. contributed human T cell CRISPR editing expertise. C.A.P. provided input throughout the study and edited the manuscript. S.C.J. supervised the GR mouse study and edited the manuscript.

## Acknowledgments

We thank the University Flow Cytometry Resource (UFCR) core facility, the Masonic Cancer Center Analytical Biochemistry core facility, and the University of Minnesota Supercomputing Institute. We are grateful to past and current members of the Kelekar laboratory for insightful discussions. This work was supported by National Institutes of Health grants R21AI148876, R03AI154388, and R01CA157971 (A.K.), R01AI38903 (S.C.J.), R01CA263090 (C.A.P.), and an MN Partnership Infrastructure Award (A.K.). T.Y. was supported by a Doctoral Dissertation Fellowship from the University of Minnesota Graduate School and W.J.V. is funded by an F32 fellowship (F32AI194558) from the NIH.

## Lead contact

Requests for further information and resources should be directed to the lead contact Dr. Ameeta Kelekar.

## References

1. Wang, R. & Green, D.R. Metabolic checkpoints in activated T cells. Nat Immunol 13, 907–915 (2012).

2. MacIver, N.J., Michalek, R.D. & Rathmell, J.C. Metabolic regulation of T lymphocytes. Annu Rev Immunol 31, 259–283 (2013).

3. Raynor, J.L., Chapman, N.M. & Chi, H. Metabolic Control of Memory T-Cell Generation and Stemness. Cold Spring Harb Perspect Biol 13 (2021).

4. van der Windt, G.J. et al. Mitochondrial respiratory capacity is a critical regulator of CD8+ T cell memory development. Immunity 36, 68–78 (2012).

5. Huang, D.C. & Strasser, A. BH3-Only proteins-essential initiators of apoptotic cell death. Cell 103, 839–842 (2000).

6. Kelekar, A. & Thompson, C.B. Bcl-2-family proteins: the role of the BH3 domain in apoptosis. Trends Cell Biol 8, 324–330 (1998).

7. Korell, F. et al. Comparative analysis of Bcl-2 family protein overexpression in CAR T cells alone and in combination with BH3 mimetics. Sci Transl Med 16, eadk7640 (2024).

8. Hijikata, M., Kato, N., Sato, T., Kagami, Y. & Shimotohno, K. Molecular cloning and characterization of a cDNA for a novel phorbol-12-myristate-13-acetate-responsive gene that is highly expressed in an adult T-cell leukemia cell line. J Virol 64, 4632–4639 (1990).

9. Hershko, T. & Ginsberg, D. Up-regulation of Bcl-2 homology 3 (BH3)-only proteins by E2F1 mediates apoptosis. J Biol Chem 279, 8627–8634 (2004).

10. Giam, M., Huang, D.C. & Bouillet, P. BH3-only proteins and their roles in programmed cell death. Oncogene 27 **Suppl 1**, S128–136 (2008).

11. Lowman, X.H. et al. The proapoptotic function of Noxa in human leukemia cells is regulated by the kinase Cdk5 and by glucose. Mol Cell 40, 823–833 (2010).

12. Karim, C.B. et al. Structural Mechanism for Regulation of Bcl-2 protein Noxa by phosphorylation. Sci Rep 5, 14557 (2015).

13. Hanse, E.A. et al. Cytosolic malate dehydrogenase activity helps support glycolysis in actively proliferating cells and cancer. Oncogene 36, 3915–3924 (2017).

14. Ploner, C., Kofler, R. & Villunger, A. Noxa: at the tip of the balance between life and death. Oncogene 27 **Suppl 1**, S84–92 (2008).

15. Fischer, S.F., Belz, G.T. & Strasser, A. BH3-only protein Puma contributes to death of antigen-specific T cells during shutdown of an immune response to acute viral infection. Proceedings of the National Academy of Sciences of the United States of America 105, 3035–3040 (2008).

16. Grayson, J.M., Weant, A.E., Holbrook, B.C. & Hildeman, D. Role of Bim in regulating CD8+ T-cell responses during chronic viral infection. J Virol 80, 8627–8638 (2006).

17. Wensveen, F.M. et al. Apoptosis threshold set by Noxa and Mcl-1 after T cell activation regulates competitive selection of high-affinity clones. Immunity 32, 754–765 (2010).

18. Dronca, R.S., et al. T cell Bim levels reflect responses to anti-PD-1 cancer therapy. JCI Insight 1 (2016).

19. Chang, C.H. et al. Posttranscriptional control of T cell effector function by aerobic glycolysis. Cell 153, 1239–1251 (2013).

20. Arlettaz, L. et al. CD45 isoform phenotypes of human T cells: CD4(+)CD45RA(-)RO(+) memory T cells re-acquire CD45RA without losing CD45RO. Eur J Immunol 29, 3987–3994 (1999).

21. McGuire, D.J. et al. Regulation of CD45 isoforms during human effector and memory CD8 T cell differentiation: Implications for T cell nomenclature. Proceedings of the National Academy of Sciences of the United States of America 122, e2322982122 (2025).

22. Michie, C.A., McLean, A., Alcock, C. & Beverley, P.C. Lifespan of human lymphocyte subsets defined by CD45 isoforms. Nature 360, 264–265 (1992).

23. Bajnok, A., Ivanova, M., Rigo, J., Jr. & Toldi, G. The Distribution of Activation Markers and Selectins on Peripheral T Lymphocytes in Preeclampsia. Mediators Inflamm 2017, 8045161 (2017).

24. Saraiva, D.P. et al. Expression of HLA-DR in Cytotoxic T Lymphocytes: A Validated Predictive Biomarker and a Potential Therapeutic Strategy in Breast Cancer. Cancers (Basel) 13 (2021).

25. O’Sullivan, D. et al. Memory CD8(+) T cells use cell-intrinsic lipolysis to support the metabolic programming necessary for development. Immunity 41, 75–88 (2014).

26. van der Windt, G.J. et al. CD8 memory T cells have a bioenergetic advantage that underlies their rapid recall ability. Proceedings of the National Academy of Sciences of the United States of America 110, 14336–14341 (2013).

27. Pearce, E.L. et al. Enhancing CD8 T-cell memory by modulating fatty acid metabolism. Nature 460, 103–107 (2009).

28. Macintyre, A.N. & Rathmell, J.C. Activated lymphocytes as a metabolic model for carcinogenesis. Cancer & metabolism 1, 5 (2013).

29. Pearce, E.L., Poffenberger, M.C., Chang, C.H. & Jones, R.G. Fueling immunity: insights into metabolism and lymphocyte function. Science 342, 1242454 (2013).

30. Chen, L. & Cui, H. Targeting Glutamine Induces Apoptosis: A Cancer Therapy Approach. Int J Mol Sci 16, 22830–22855 (2015).

31. Jewell, J.L. et al. Metabolism. Differential regulation of mTORC1 by leucine and glutamine. Science 347, 194–198 (2015).

32. Nicklin, P. et al. Bidirectional transport of amino acids regulates mTOR and autophagy. Cell 136, 521–534 (2009).

33. Magnuson, B., Ekim, B. & Fingar, D.C. Regulation and function of ribosomal protein S6 kinase (S6K) within mTOR signalling networks. Biochem J 441, 1–21 (2012).

34. Qin, X., Jiang, B. & Zhang, Y. 4E-BP1, a multifactor regulated multifunctional protein. Cell Cycle 15, 781–786 (2016).

35. Pollizzi, K.N. et al. mTORC1 and mTORC2 selectively regulate CD8(+) T cell differentiation. J Clin Invest 125, 2090–2108 (2015).

36. Takahara, T., Amemiya, Y., Sugiyama, R., Maki, M. & Shibata, H. Amino acid-dependent control of mTORC1 signaling: a variety of regulatory modes. J Biomed Sci 27, 87 (2020).

37. Beyer, S.R. et al. Identification of cysteine residues in human cationic amino acid transporter hCAT-2A that are targets for inhibition by N-ethylmaleimide. J Biol Chem 288, 30411–30419 (2013).

38. Luo, Z. et al. Co-delivery of 2-Deoxyglucose and a glutamine metabolism inhibitor V9302 via a prodrug micellar formulation for synergistic targeting of metabolism in cancer. Acta Biomater 105, 239–252 (2020).

39. Okunushi, K. et al. JPH203, a newly developed anti-cancer drug, shows a preincubation inhibitory effect on L-type amino acid transporter 1 function. J Pharmacol Sci 144, 16–22 (2020).

40. Shrestha, S. et al. Tsc1 promotes the differentiation of memory CD8+ T cells via orchestrating the transcriptional and metabolic programs. Proceedings of the National Academy of Sciences of the United States of America 111, 14858–14863 (2014).

41. Yang, K. et al. T cell exit from quiescence and differentiation into Th2 cells depend on Raptor-mTORC1-mediated metabolic reprogramming. Immunity 39, 1043–1056 (2013).

42. Owen, O.E., Kalhan, S.C. & Hanson, R.W. The key role of anaplerosis and cataplerosis for citric acid cycle function. J Biol Chem 277, 30409–30412 (2002).

43. Raud, B. et al. Etomoxir Actions on Regulatory and Memory T Cells Are Independent of Cpt1a-Mediated Fatty Acid Oxidation. Cell Metab 28, 504–515.e507 (2018).

44. Araki, K. et al. mTOR regulates memory CD8 T-cell differentiation. Nature 460, 108–112 (2009).

45. Sengupta, S., Peterson, T.R. & Sabatini, D.M. Regulation of the mTOR complex 1 pathway by nutrients, growth factors, and stress. Mol Cell 40, 310–322 (2010).

46. Wang, R. et al. The transcription factor Myc controls metabolic reprogramming upon T lymphocyte activation. Immunity 35, 871–882 (2011).

47. Sandalova, E., Wei, C.H., Masucci, M.G. & Levitsky, V. Regulation of expression of Bcl-2 protein family member Bim by T cell receptor triggering. Proceedings of the National Academy of Sciences of the United States of America 101, 3011–3016 (2004).

48. Gubser, P.M. et al. Aerobic glycolysis but not GLS1-dependent glutamine metabolism is critical for anti-tumor immunity and response to checkpoint inhibition. Cell Rep 43, 114632 (2024).

49. Nabe, S. et al. Reinforce the antitumor activity of CD8(+) T cells via glutamine restriction. Cancer Sci 109, 3737–3750 (2018).

50. Wensveen, F.M. et al. Pro-apoptotic protein Noxa regulates memory T cell population size and protects against lethal immunopathology. J Immunol 190, 1180–1191 (2013).

51. Ganuza, M. et al. Neurobeachin regulates hematopoietic progenitor differentiation and survival by modulating Notch activity. Blood Adv 8, 4129–4143 (2024).

52. Huang, L., Wei, B., Zhao, Y., Gong, X. & Chen, L. DYNLT1 promotes mitochondrial metabolism to fuel breast cancer development by inhibiting ubiquitination degradation of VDAC1. Mol Med 29, 72 (2023).

53. Huang, Y. et al. The cationic amino acid transporters CAT1 and CAT3 mediate NMDA receptor activation-dependent changes in elaboration of neuronal processes via the mammalian target of rapamycin mTOR pathway. J Neurosci 27, 449–458 (2007).

54. Yang, Y., Hu, Y.H. & Liu, Y. Wdfy1 deficiency impairs Tlr3-mediated immune responses in vivo. Cell Mol Immunol 17, 1014–1016 (2020).

55. Cheng, Z.Y., He, T.T., Gao, X.M., Zhao, Y. & Wang, J. ZBTB Transcription Factors: Key Regulators of the Development, Differentiation and Effector Function of T Cells. Front Immunol 12, 713294 (2021).

56. Morsi, R.Z., Hage-Sleiman, R., Kobeissy, H. & Dbaibo, G. Noxa: Role in Cancer Pathogenesis and Treatment. Curr Cancer Drug Targets 18, 914–928 (2018).

57. Bajpai, R. et al. Targeting glutamine metabolism in multiple myeloma enhances BIM binding to BCL-2 eliciting synthetic lethality to venetoclax. Oncogene 35, 3955–3964 (2016).

58. Yan, Y. et al. Phosphatase PHLPP2 regulates the cellular response to metabolic stress through AMPK. Cell Death Dis 12, 904 (2021).

59. Webber, B.R., et al. Highly efficient multiplex human T cell engineering without double-strand breaks using Cas9 base editors. Nat Commun 10, 5222 (2019).

60. Joshua, B., Thomas, K., Adam, H. & Ying, Z. (2019).

61. Kim, D., Langmead, B. & Salzberg, S.L. HISAT: a fast spliced aligner with low memory requirements. Nat Methods 12, 357–360 (2015).

62. Liao, Y., Smyth, G.K. & Shi, W. featureCounts: an efficient general purpose program for assigning sequence reads to genomic features. Bioinformatics 30, 923–930 (2014).

63. Team, R.C. (R Foundation for Statistical Computing; 2024).

64. McCarthy, D.J., Chen, Y. & Smyth, G.K. Differential expression analysis of multifactor RNA-Seq experiments with respect to biological variation. Nucleic Acids Res 40, 4288–4297 (2012).

65. Yu, G., Wang, L.G., Han, Y. & He, Q.Y. clusterProfiler: an R package for comparing biological themes among gene clusters. OMICS 16, 284–287 (2012).

66. Wickham, H. ggplot2: Elegant Graphics for Data Analysis. (Springer-Verlag, 2016).

67. Kolde, R. Pheatmap: pretty heatmaps. R package version 1, 726 (2019).

