## Supplemental Figure1 for "Bcl-2 protein Noxa is required for metabolic reprogramming to glutamine dependence and for apoptosis in stimulated human CD8^+^ T cells"

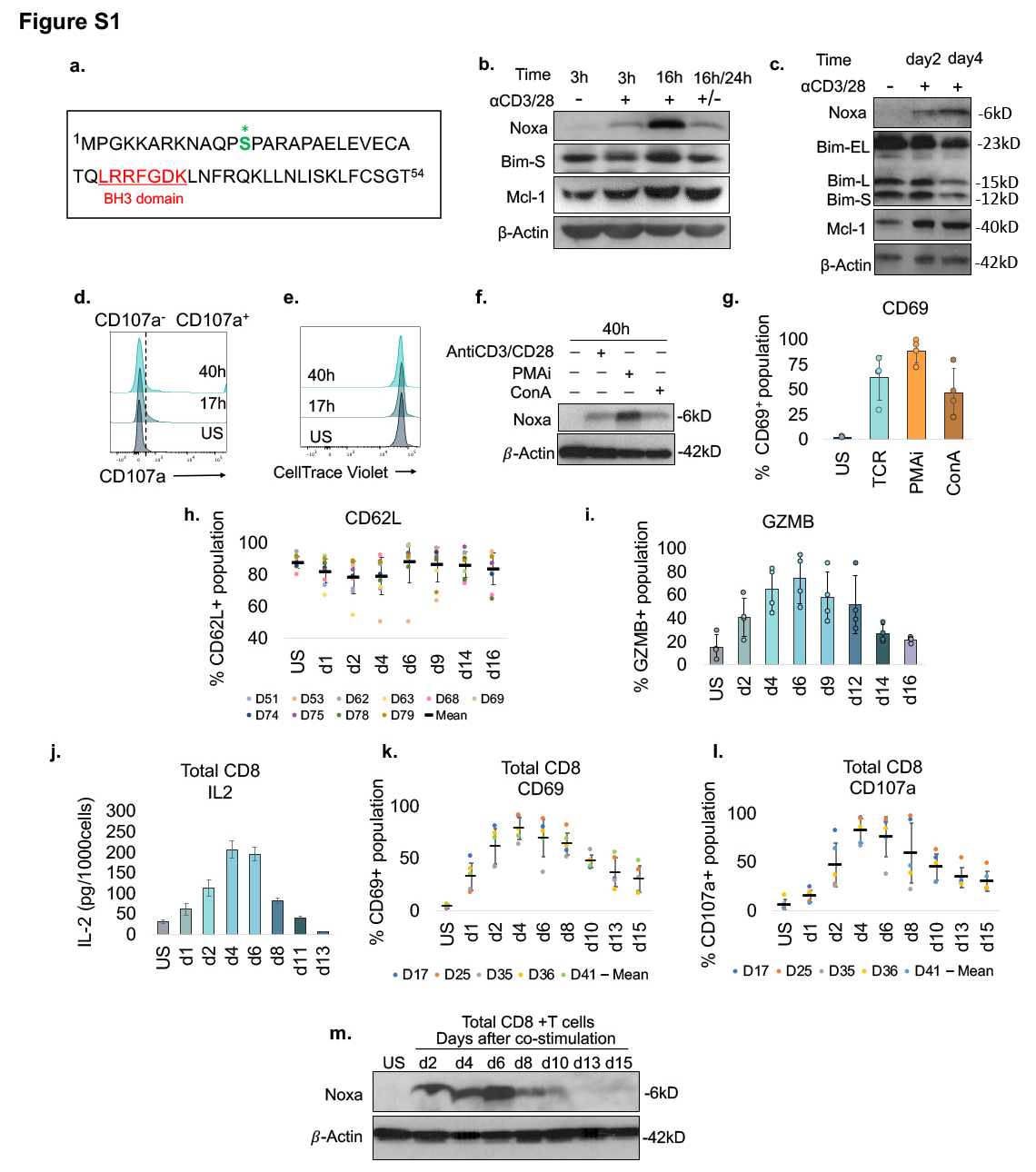

**Figure S1. Noxa protein is induced upon activation and remains expressed during proliferation and differentiation of naïve and total CD8⁺ human T cells. (Related to Figure 1). a.** Sequence of 54-residue human Noxa showing the BH3 domain residues (in red) and the phosphorylated serine (in green). **b-c.** Western blot analysis of Bim and Mcl-1 protein expression in US, 3h and 16h stimulated and 16+/24h- S/US pan T cells (**b**); and US, day 2 and day 4-stimulated naïve CD8^+^ T cells (**c**). **d-e.** Representative histograms showing CD107a (**d**) and Cell Trace Violet, CTV (**e**) levels in unstimulated, 17h and 40h stimulated naïve CD8^+^ T cells. **f-g**. Noxa protein levels (**f**) in naïve CD8⁺ T cells left unstimulated or stimulated for 40h with anti-CD3/CD28, PMA/ionomycin (PMAi) or Concanavalin A (ConA), and CD69 induction following these treatments (**g**) averaged from 4 independent donors. **h-i.** CD62L (**h**) and Granzyme B (GZMB) (**i**) expression in naive CD8^+^ T cells from 10 individual donors at the indicated times after stimulation (see Fig **1i, j**). **j-l.** Total CD8+ T cells from multiple donors were analyzed at the indicated times after stimulation for IL-2 production (**j**), CD69(**k**) and CD107a (**l**). Data are representative of 4-10 independent experiments. **m.** Western blot showing Noxa levels in total CD8+ T cells co-stimulated at the indicated times in vitro.
