## Supplemental Figure2 for "Bcl-2 protein Noxa is required for metabolic reprogramming to glutamine dependence and for apoptosis in stimulated human CD8^+^ T cells"

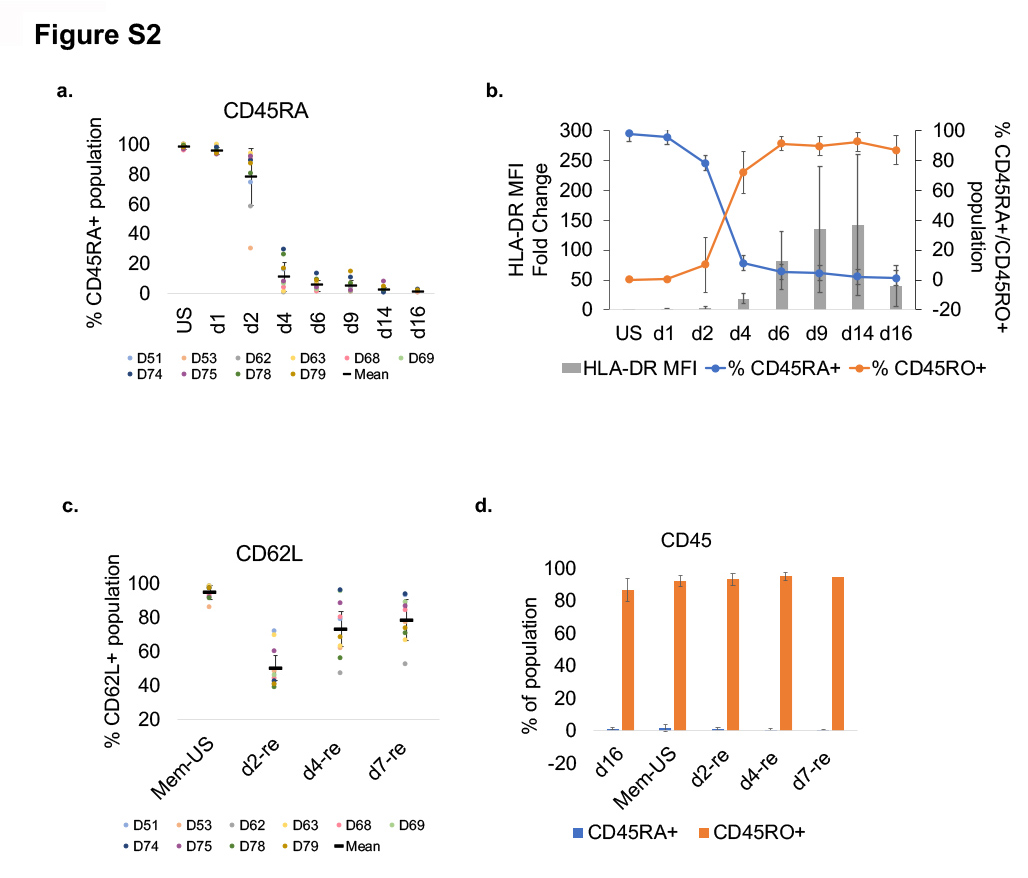

**Figure S2. Detection and re-stimulation of memory-like CD8^+^ T cells. (Related to Figure 2). a**. Naïve CD8^+^ T cells were analyzed at the indicated time points after stimulation for CD45RA positivity (data from 10 independent experiments). **b.** Flow cytometric analysis of CD45RA⁺ and CD45RO⁺ populations and HLA-DR mean fluorescence intensity fold change in naïve CD8⁺ T cells at the indicated time points after stimulation (n = 10). **c, d.** Memory populations were re-stimulated and analyzed for CD62L (**c**) and CD45 (**d**) positivity. Data are representative of 4-10 independent experiments. (D=donor).
