## Supplemental Figure3 for "Bcl-2 protein Noxa is required for metabolic reprogramming to glutamine dependence and for apoptosis in stimulated human CD8^+^ T cells"

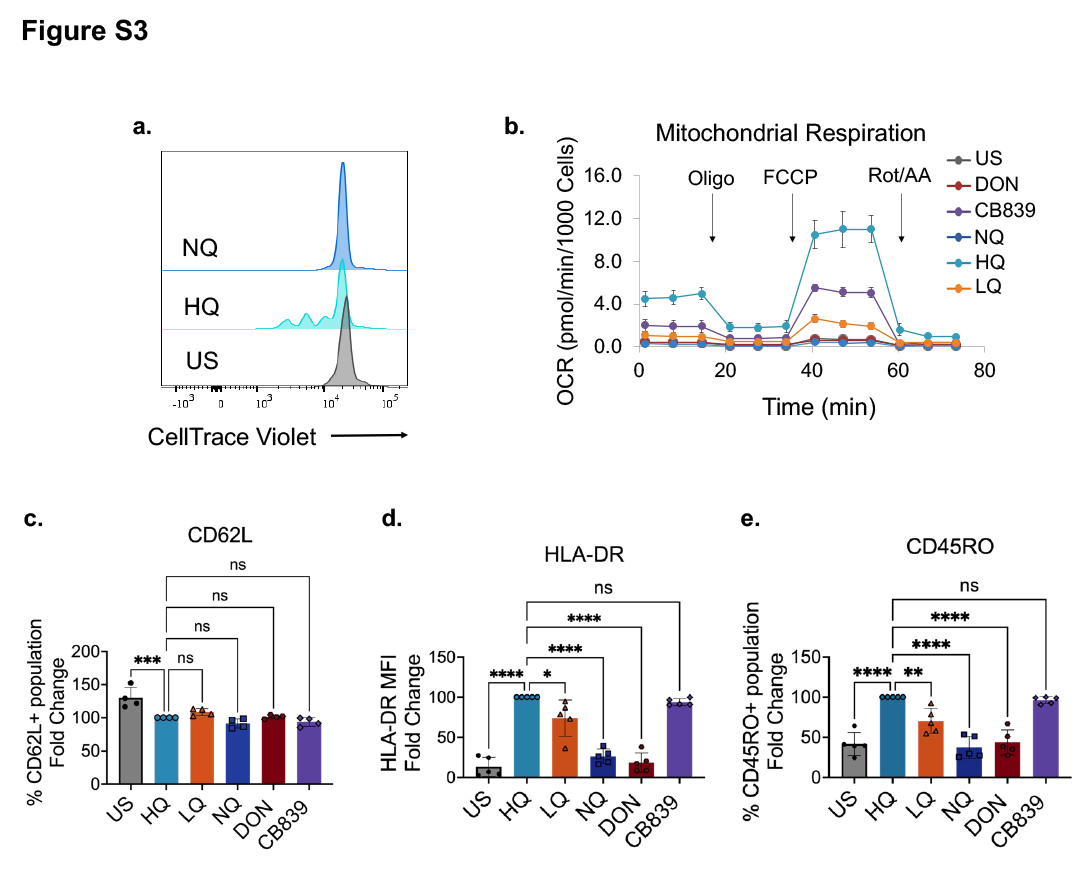

**Figure S3. Glutamine is required for metabolic fitness, late activation and differentiation in stimulated CD8^+^ T cells. (Related to Figure 3). a**. Proliferation rate in CTV labelled naïve CD8^+^ T cells 48 hours after stimulation with anti-CD3/CD28 antibodies in high glutamine (HQ) or glutamine-free (NQ) medium. **b**. OCR in unstimulated (US) and 40-hour post-stimulated representative donor CD8⁺ T cells cultured in HQ, LQ, or NQ, in the presence or absence of CB839 (2uM) or DON (200nM). **c-f**. Analyses of CD8⁺ T cell response to varying glutamine conditions and CB839 and DON treatments 2 or 4 days, post-stimulation. Readouts include percent CD62L⁺ cells (**c**) and cell division (**d**) on day 2, and percent CD45RO⁺ cells (**e**), and HLA-DR MFI (**f**) on day 4 post-stimulation (n = 8).
