## Supplemental Figure4 for "Bcl-2 protein Noxa is required for metabolic reprogramming to glutamine dependence and for apoptosis in stimulated human CD8^+^ T cells"

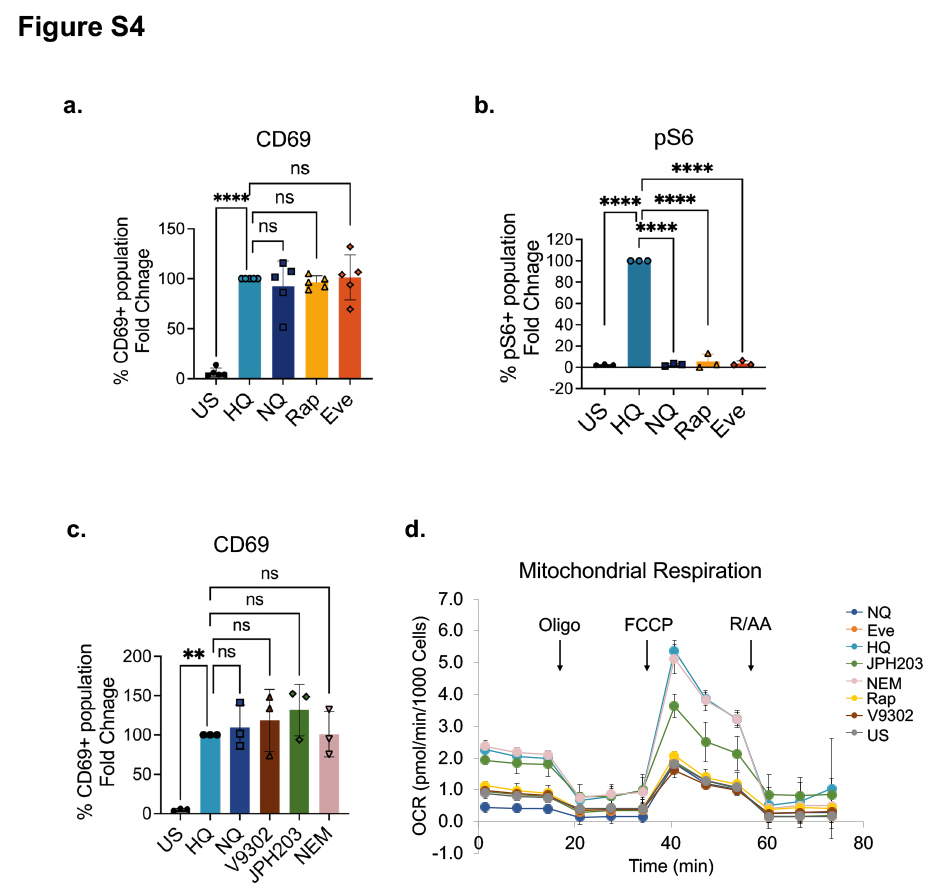

**Figure S4. Both glutamine deprivation and mTOR inhibition have no effect on CD8⁺ T cell activation but diminish mitochondrial respiratory capacity. (Related to Figure 4). a, b.** Analysis of CD8^+^ T cell responses under mTORC1 inhibition. Fold change in CD69 (**a**) on day 2 and phospho-S6 (**b**) positivity on day4 of stimulation, relative to HQ group. n = 3). **c.** Fold change in CD69 levels in CD8⁺ T cells treated with indicated amino acid transporter inhibitors, relative to positive (HQ) control (n = 3). **d.** Oxygen consumption rate (OCR) of unstimulated (US) and 40-hour post-stimulated CD8⁺ T cells, from a representative donor, cultured with or without glutamine, or in the presence of rapamycin, everolimus, V-9302, JPH203, or NEM, as indicated (n=5).
