## Supplemental Figure5 for "Bcl-2 protein Noxa is required for metabolic reprogramming to glutamine dependence and for apoptosis in stimulated human CD8^+^ T cells"

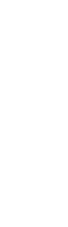

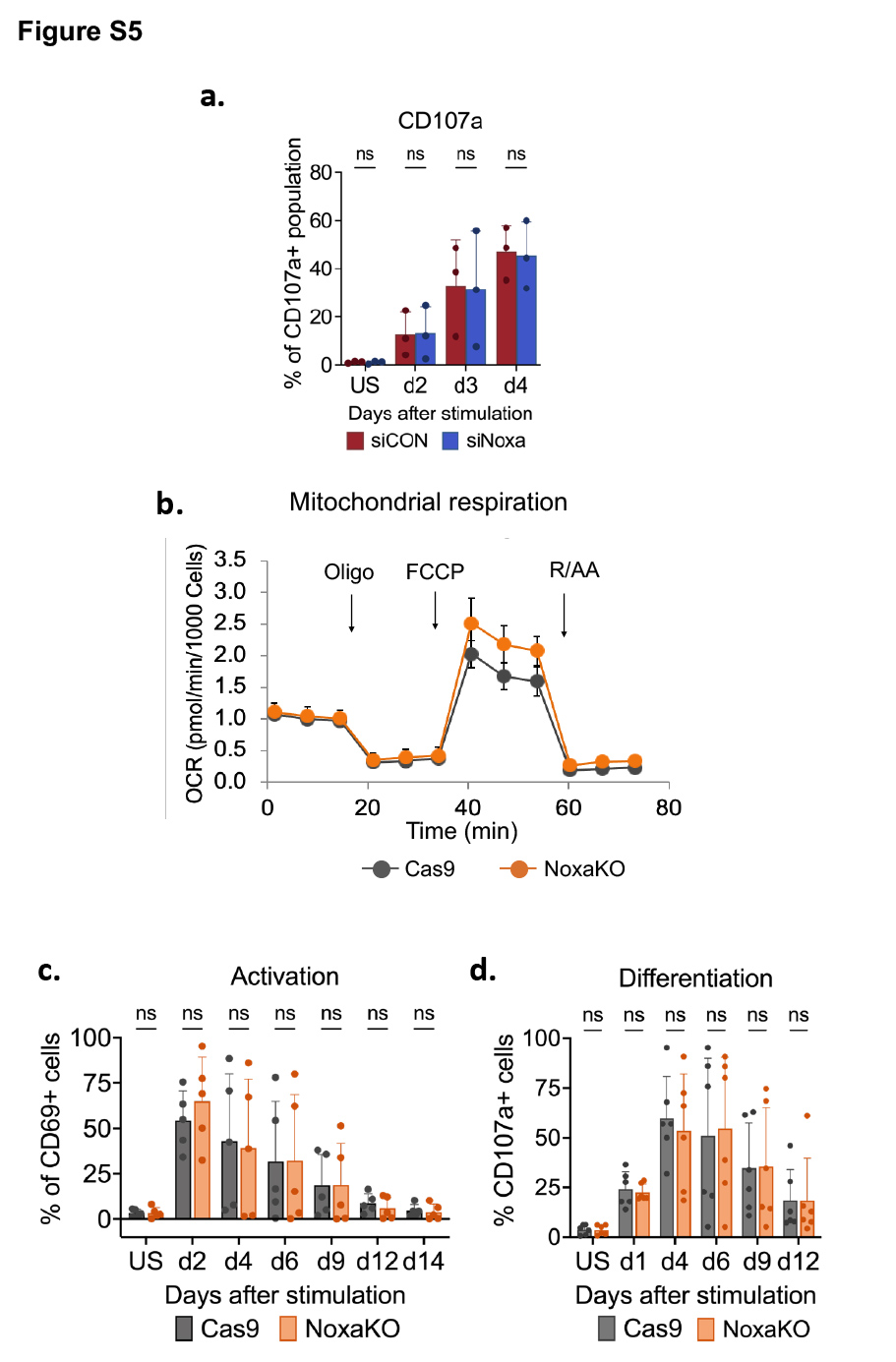

**Figure S5.** **Noxa knock down or loss does not affect activation, mitochondrial respiration, or differentiation of CD8⁺ T cells. (Related to Figure 5). a.** Percentage of CD107a⁺ cells in siCON- and siNoxa-treated CD8⁺ T cells at indicated time points following stimulation (n = 3). **b.** OCR of Cas9-control and NoxaKO CD8+ T cells from a representative donor, 48h post-stimulation (n=5). **c. d.** Analysis of Cas9-control and NoxaKO CD8^+^ T cells for CD69 (**c**) and CD107a (**d**) expression at indicated times post-stimulation (n=5-6).
