## Supplemental Figure6 for "Bcl-2 protein Noxa is required for metabolic reprogramming to glutamine dependence and for apoptosis in stimulated human CD8^+^ T cells"

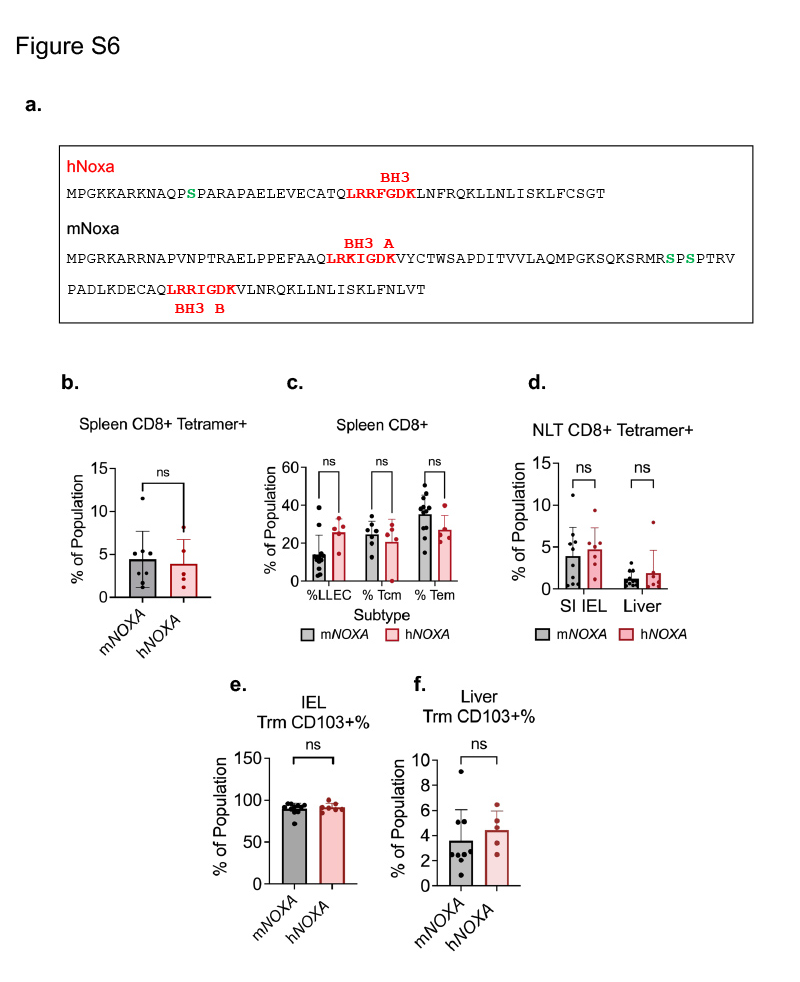

**Figure S6. CD8⁺ T cells from gene replacement mice expressing h*NOXA* show normal differentiation after LCMV infection. (Related to Figure 6). a.** Protein sequence of human Noxa (hNoxa) and mouse Noxa (mNoxa) showing BH3 domains in red and relevant serine residues (amino terminal in hNoxa and internal in mNoxa) in green. **b-f.** m*NOXA* and h*NOXA* mice were infected with lymphocytic choriomeningitis virus (LCMV) and analyzed 12 weeks post-infection. Spleen, liver, and small intestine intraepithelial lymphocytes (SI IELs) were isolated and analyzed by flow cytometry on live, unfixed cells **(b).** Percent CD8⁺ tetramer⁺ cells and specific CD8⁺ T cell subsets in the spleen **(c). d–f.** Percent CD8⁺ tetramer⁺ cells and tissue-resident memory T cells (Trm) in non-lymphoid tissues, including SI IELs and liver (n = 5-12).
