## Supplemental Figure7 for "Bcl-2 protein Noxa is required for metabolic reprogramming to glutamine dependence and for apoptosis in stimulated human CD8^+^ T cells"

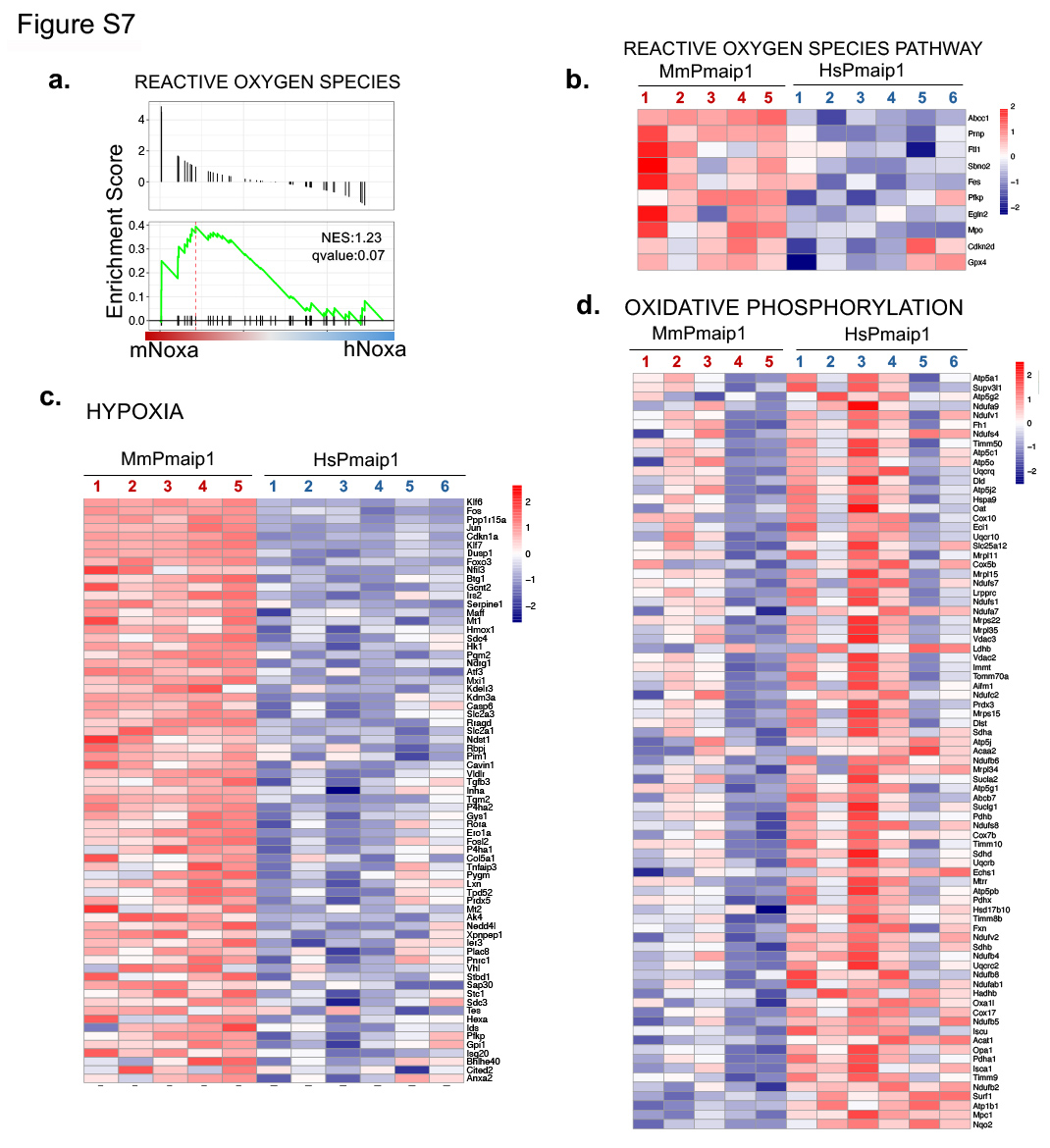

**Figure S7. h*NOXA*-expressing CD8⁺ T cells display reduced stress-related and increased growth-related early gene signatures. (Related to Figure 7). a.** GSEA enrichment plot of the Hallmark *Reactive Oxygen Species* pathway comparing 48-hour-stimulated m*NOXA* and h*NOXA* CD8⁺ T cells. NES and nominal P values are shown. **b–d.** Heatmaps depicting transcript abundance of top leading-edge genes within the Hallmark gene sets *Reactive Oxygen Species* (n = 10), *Hypoxia* (n = 69), and *Oxidative Phosphorylation* (n = 81), where n indicates the number of leading-edge genes.
